# Quantitative extrapolation from single-tags (QuEST) immunofluorescence microscopy to derive TCR signalosome stoichiometries in human primary T cells

**DOI:** 10.64898/2026.03.28.715001

**Authors:** Panyu Fei, Michael L. Dustin

## Abstract

Upon T cell receptor (TCR) engagement, a T cell forms an immunological synapse (IS) with an antigen-presenting cell (APC), which can be mimicked by purified ligands on supported lipid bilayers (SLBs)^1,2^. Microvilli actively scan the surface; upon initial engagement, F-actin-dependent TCR microclusters form, and the central supramolecular activation cluster (cSMAC) sustains TCR signaling in CD8⁺ T cells^3,4^. Although signaling activities within the IS have been observed qualitatively through total internal reflection immunofluorescence microscopy^5–7^, the stoichiometric relationships among the components of the TCR signalosome remain unknown. In this study, we employed a two-step approach to quantify the components of the TCR signalosome. First, Jurkat cell lines expressing GFP-tagged proteins on a knockout background were used to calibrate fluorescence intensity (IF) signals against molecular copy numbers, based on measurements of single-tag signals and multiple corrections. In the second step, this calibration was applied to determine the stoichiometries of key TCR signalosome components, including TCR, CD8, CD28, CD45, PD-1, Lck, ZAP-70, LAT, and PLCγ1, across scanning, early activation, and sustained activation states in human primary T cells. We refer to the method as quantitative extrapolation from single-tags (QuEST) immunofluorescence microscopy. Applying the QuEST, we were surprised to find that the ZAP-70:TCR ratio in microclusters and the cSMAC was 1:1, far from the potential 10:1 ratio. Nanoscale structures of the TCR signalosome were further captured using direct stochastic optical reconstruction microscopy (dSTORM), confirming that ZAP-70 was strongly co-localized with the TCR. Moreover, we applied QuEST to confirm the presence of T cell intrinsic CD28 recruitment, independent of CD80 or CD86 on SLBs, during TCR activation. This T cell intrinsic CD28 recruitment could be disrupted through engagement of PD-1 with PD-L1 on SLBs. This shows that PD-1 engagement can disrupt T cell intrinsic CD28 costimulation. QuEST provides a broadly applicable pipeline for quantitative analysis of TCR signalosomes in human primary cells, enabling a quantitative basis for the rational manipulation and engineering of the TCR signalosome in immunotherapies.

## Introduction

T cells are essential for immune homeostasis, protecting the host from pathogens, eliminating tumor cells, and suppressing pathological immune responses^4,8,9^. Central to these processes is the T cell antigen receptor (TCR), a covalent heterodimer generated by somatic recombination and non-covalently associated with two CD3 heterodimers (CD3εγ and CD3εδ) and the CD247 homodimer (ζζ). This multi-chain complex confers the exquisite specificity and sensitivity required for antigen recognition by T cells^10^. Upon engagement of peptide–major histocompatibility complex (pMHC) on an antigen-presenting cell, numerous surface and intracellular signaling molecules of CD8^+^ T cells, including the coreceptor CD8, phosphatase CD45, adapter LAT, kinases Lck and ZAP-70, and lipase PLCγ1, undergo pronounced spatial and temporal reorganizations. The TCR contains 10 immunotyrosine activation motifs (ITAMs), which can each bind ZAP-70. A single pMHC engagement event can recruit up to six ZAP-70 molecules, but the number of TCRs involved was not determined^6^, precluding the determination of the stoichiometry. The presence of 10 ITAMs in the TCR is functionally important, but the main impact of reducing ITAM number is to eliminate the binding of additional signaling molecules like Vav and SHP1^11,12^. The outcome of TCR signaling is further modified by costimulatory receptors, such as CD28, and inhibitory receptors, such as PD-1. These components are recruited to, or excluded from, sites of TCR clustering to form the nanoscale architecture of the TCR signalosome, comprising pre-engagement scanning, early microcluster formation, followed by sustained signaling within the cSMAC in the mature IS^4,13^. TCR signalosome function is impaired in primary immunodeficiencies lacking or significantly modifying signalosome components and may be quantitatively altered by coding polymorphisms associated with altered disease risk.

Despite its central role in T cell activation, the numbers and stoichiometric relationships among the signaling molecules within the TCR signalosome are not known. Bulk analysis using tetramer stimulation and mass spectrometry has provided guidance^14^, but this is hard to relate back to physiological activation in an immunological synapse. The lack of quantitative insight limits our capacity to anticipate how genetic variation or synthetic manipulation, such as PD-1 or CD28 blockade or agonism, or TCR engineering, reshapes signaling outcomes. These knowledge gaps contribute to challenges in predicting the impact of genetic variants and the optimization of immunotherapies^15^. An accurate blueprint of signalosome nanoarchitecture would provide a mechanistic foundation for rationally engineering TCR-based therapeutics. Addressing these fundamental and translational needs requires determination of the stoichiometry of the TCR signalosome, an effort that demands quantification of the numbers and ratios of molecular signalosome components at nanometer resolution in the crowded and heterogeneous cellular regions.

Over recent decades, the single-molecule imaging and force strategies^16–19^ have transformed our ability to visualize and interrogate individual molecules with exceptional sensitivity. However, these approaches break down in dense, crowded cellular environments. Previous attempts, including intensity-to-density conversion using fluorescent standard beads^20,21^, particle-to-molecule mapping via electron microscopy^22,23^, microfluidic antibody-counting approaches^24^, and stepwise photobleaching analyses^25^, have provided valuable insights but are constrained by indirect calculation, low throughput, specialized instrumentation, and limitations in biological complexity. Furthermore, immunofluorescence microscopy involves multiple steps with less than 100% efficiency, and appreciating the cumulative effect of these factors is non-trivial. Even methods such as quantitative points accumulation in nanoscale topography (qPAINT)^26^ that enable signal molecule counting still require specialized reagents and a long imaging process, and need more accurate quantification of molecular loss and antibody-targeting efficiency. The principle of relating fluorescence intensities to single-molecule intensities, which enables extrapolation to molecular densities, has recently been applied to molecular estimation^27,28^, but the assay has never been carefully calibrated under various imaging conditions in crowded and complex biological samples. Therefore, new methods are required for careful fluorescence calibration to tackle problems such as TCR signalosome stoichiometries.

Here, we have developed a methodological framework that overcomes the above barriers by enabling direct and accurate estimation of molecular densities and stoichiometric relationships from TIRFM by extending the single-molecule paradigm to crowded biological samples with multi-faceted corrections, which we refer to as **Qu**antitative **E**xtrapolation from **S**ingle-**T**ags (**QuEST**) immunofluorescence microscopy. In addition to the single-tag-based calibration, we have anticipated sources of systematic error impacting the overall accuracy of the pipeline, including illumination intensity, exposure time, temperature, stability, fixation, self-quenching, fluorescence resonance energy transfer (FRET), 2D illumination uniformity, molecular loss during sample treatments, imaging distance from the substrate, and measuring area. These factors are fully corrected by matching the calibration and experimental acquisition conditions within the experimentally determined linear range. We have used images of knockout Jurkat cell lines expressing GFP-tagged versions of key signaling proteins to follow their fate from live imaging through fixation, permeabilization, staining, washing, and microscopy imaging. The cumulative losses of the GFP signal and how this relates to the immunofluorescence signature can then be followed to determine target-specific correction factors to go from the IF intensities to molecular densities in an environment that is very similar to primary cell subjects. Using QuEST, we accurately estimate the number, density, and stoichiometries of TCR signalosome components across scanning (bulk and pre-cluster), early microcluster, and sustained cSMAC IS phases in human primary T cells. A surprising finding was that the stoichiometric ratio of the key kinase ZAP-70 to TCR remains tightly constrained at approximately 1.0 across all conditions, including scanning, despite the presence of up to 10 potential ZAP-70 binding sites on the TCR. The local density of TCR and levels of ZAP-70 phosphorylation display a greater range across conditions, suggesting that these are important parameters determining T cell response. The near unity stoichiometry of TCR:ZAP-70 has implications for how the TCR functions, leaving binding sites open for other SH2-expressing signaling proteins.

## Results

### Quantification of single-tag fluorescence

The copy number and surface density of protein molecules, as well as their stoichiometric relationships within large complexes, are fundamental to understanding their behavior and function. Yet application of single-molecule fluorescence-based calibration remains challenging due to the sheer abundance, nanoscale dimensions, and mobility of proteins in biological samples. To overcome these limitations, we extended the single-molecule paradigm using data acquired by TIRFM and sCMOS sensors that can determine the fluorescence of a single-molecule tag (a purified ligand, a GFP-fused molecule, or an antibody complex) and the same tag used in a biological setting at high density. QuEST starts with the principle of single-molecule fluorescence microscopy by determining the fluorescence intensity of an isolated tag (I_S_) under the same conditions used to measure the total fluorescence from multiple molecule tags (I_M_), the total number of molecules (N) can be calculated as N=I_M_/I_S_. Dividing N by the measured area (A) yields the molecular density (Fig. 1a and Methods). As this principle has been applied previously to the calibration of ligands on the SLBs^27^, we started with the careful quantification of single tags under various conditions and the calibration of SLB samples as an initial step towards application to more complex biological samples. The first challenge relates to the accurate assessment of single-tag fluorescence, so that it is reproducible from day to day.

**Fig. 1.**
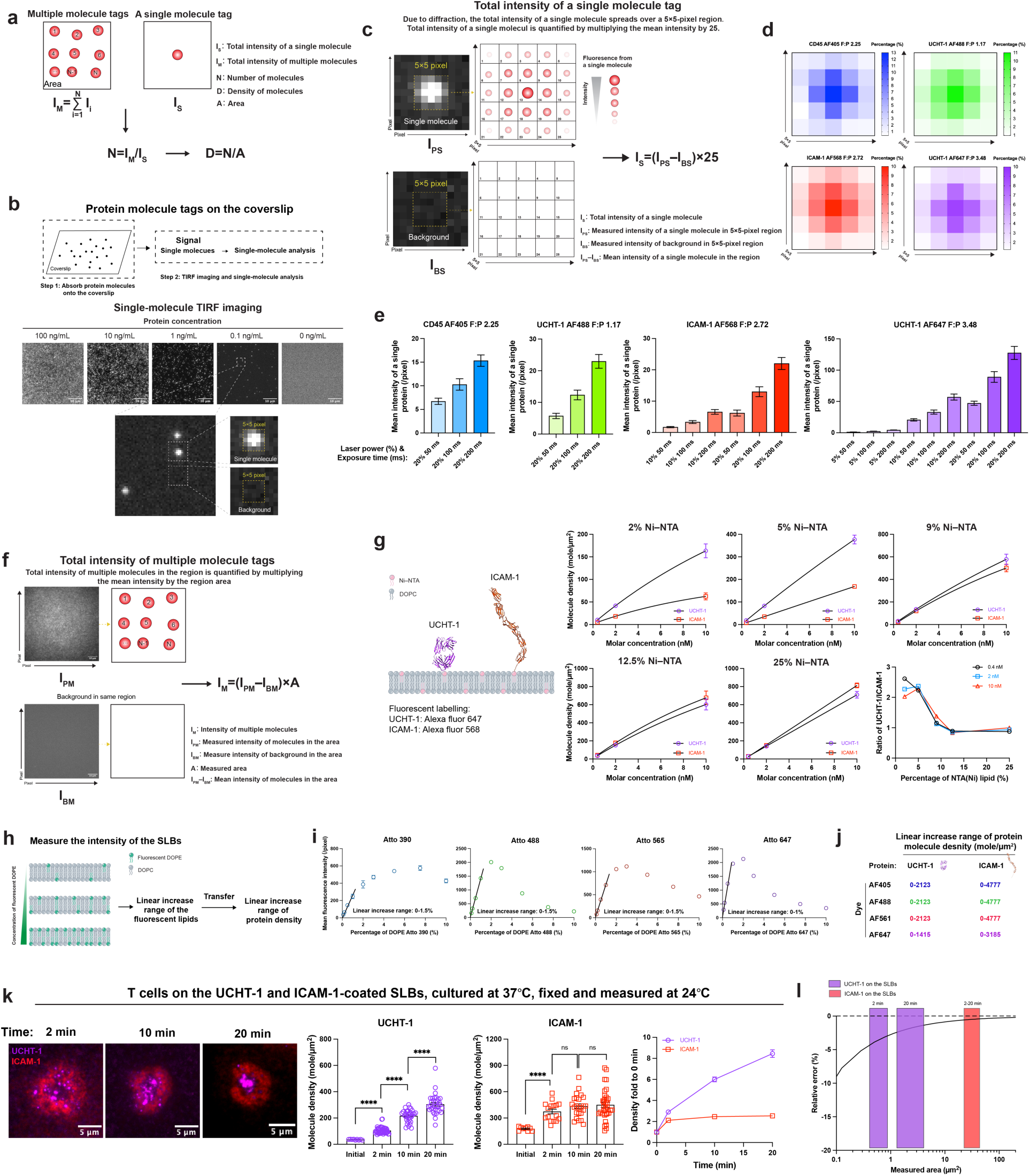
Principle of the QuEST method and quantification of molecule numbers and densities. **a**, Schematic illustrating the principle of QuEST for quantifying molecule numbers and surface densities. **b**, Isolation and detection of fluorescence-labelled single molecules by TIRFM. **c**, Workflow for quantifying single-molecule fluorescence intensity. **d**,**e**, Intensity distribution within a 5×5-pixel region (**d**) and corresponding mean fluorescence intensity (**e**) of single CD45 AF405, UCHT-1 AF488, ICAM-1 AF568, and UCHT-1 AF647 molecules. **f**, Workflow for quantifying fluorescence intensity of multiple molecules. **g**, Quantification of UCHT-1 and ICAM-1 densities on supported lipid bilayers (SLBs) containing different percentages of Ni–NTA using QuEST. **h**,**i**, Schematic (**h**) and quantification results (**i**) determining the linear dynamic range of DOPE with different fluorescence levels. **j**, Linear density ranges of UCHT-1 and ICAM-1 under different fluorescence conditions. **k**, T cells clustering UCHT-1 and ICAM-1 on SLBs, with dynamic molecular densities quantified. **l**, Relative errors in quantifying UCHT-1 and ICAM-1 densities using QuEST. Data in **g** and **i** are presented as mean ± SD; data in **e** and **k** are presented as mean ± SEM. Statistical significance was assessed using an unpaired two-tailed Student’s *t*-test (\*\*\*\**P* ≤ 0.0001; ns, not significant). In **e**, data were obtained from 16 single molecules. In **g** and **i**, data were obtained from three independent TIRF images. The structure of UCHT-1 was derived from PDB entry 3FZU, and the structure of ICAM-1 was generated using AlphaFold3 from the corresponding protein sequence.

Operationally, we diluted fluorescently labelled tags to obtain sparse, well-isolated signals on a glass coverslip and recorded their signals using TIRFM. This enabled precise measurement of individual tag intensities and local background, free from interference by neighboring emitters (Fig. 1b,c). Single-molecule identity was confirmed by stepwise photobleaching consistent with the fluorophore to protein tag (F:P) ratio of fluorescently labelled protein or the monomeric property of fluorescent proteins (Extended Data Fig. 1a). Using a 100×, 1.5 NA objective and an sCMOS camera with 11-µm pixel size, fluorescence from each tag was fully captured within a 5×5-pixel region, consistent with the diffraction limit (Fig. 1c,d, and Methods). We verified that single-tag intensity depends on excitation power, camera exposure time, fluorophore type, and dye valency (Fig. 1e and Extended Data Fig. 1b). To broaden applicability, we assessed additional variables affecting single-tag intensity. Proteins imaged immediately after thawing (“fresh”) and the same tags stored for two months at 4 °C (“old”) yielded indistinguishable single-spot intensities under identical conditions, demonstrating stability of both reagents and instrumentation over extended periods (Extended Data Fig. 1c), although this may vary by tag or tag type (protein, antibody, etc). We further evaluated temperature dependence, as fixed samples are imaged at room temperature, whereas live cell imaging of T cells is typically performed at 37 °C. Across multiple proteins labelled with four different fluorophores at varying F:P ratios, as well as distinct protein species, mean fluorescence intensity was consistently lower at 37 °C than at 24 °C (Extended Data Fig. 1d–f), indicating that temperature-specific I_S_ values are required for QuEST. Paraformaldehyde (PFA) fixation did not alter intensity for organic dye-labelled tags under different imaging conditions (Extended Data Fig. 1g,h), but GFP exhibited ∼40% intensity loss after fixation, likely due to partial denaturation (Extended Data Fig. 1i), necessitating the use of fixation-adjusted I_S_ values for GFP-tagged molecules. We propose that this should always be investigated for fluorescence proteins when comparing live and fixed samples.

Unlike single-tag isolation and quantification (Fig. 1b–e), in regions containing dense tags, the total fluorescence is obtained by multiplying the mean molecular intensity by the measured area (Fig. 1f and Methods). Conceptually, QuEST therefore enables direct, unconverted, and in situ quantification across sparse and dense regimes, and across static (e.g., coverslip) and fluidic (e.g., cell membranes, SLBs) surfaces within the range of TIRFM. Accurate measurement requires that all pixel intensities remain below the camera’s saturation limit (4095 for 12-bit, 65535 for 16-bit modes) by adjusting imaging conditions (laser power or camera exposure time). Finally, we showed the relative error in using the QuEST method in quantification (Extended Data Fig. 2a and Methods), and determined that accuracy depends on the measurement area, with larger regions yielding reduced error, as expected (Extended Data Fig. 2b).

### Calibration of ligands on SLBs

The theoretical foundation of QuEST prompted us to experimentally validate its performance by directly quantifying molecular copy numbers and surface densities on SLBs. We first applied the method to determine the densities of UCHT-1 (Fab fragment throughout the study; sequence in Methods) and ICAM-1 (extracellular domain throughout the study; sequence in Methods) on SLBs. Because the SLBs have a well-established thickness of ∼5 nm^29^, we began by addressing whether axial variations in TIRF illumination might influence intensity measurements. Using UCHT-1 AF488 as an example, we compared mean single-molecule intensities for proteins incorporated onto SLBs versus those adsorbed directly onto a coverslip. For SLB measurements, we prepared SLBs containing a low fraction of UCHT-1 AF488 diluted 1:2000 into UCHT-1 AF647 (total UCHT-1 density: 36 molecules/µm^2^) together with ICAM-1 (177 molecules/µm^2^). Upon engagement with the SLBs, CD8⁺ T cell blasts (hereafter “T cells”) clustered UCHT-1 into the central region of the interface, allowing us to resolve isolated single AF488-labelled UCHT-1 molecules from abundant AF647-labelled UCHT-1 molecules in the cSMAC (Extended Data Fig. 3a). For coverslip-bound molecules, UCHT-1 AF488 was directly immobilized and quantified (Extended Data Fig. 3b). The mean fluorescence intensities of single UCHT-1 AF488 molecules on SLBs and coverslip were indistinguishable (Extended Data Fig. 3c), indicating that TIRFM illumination is effectively uniform across the ∼5-nm axial range, and that intensities measured on coverslip can be directly applied to SLB conditions.

Using QuEST, we then quantified UCHT-1 and ICAM-1 densities on SLBs under varying imaging conditions and obtained highly consistent results (Extended Data Fig. 4a), demonstrating the stability and accuracy of the method. At low Ni–NTA concentrations with fewer binding sites, UCHT-1 bound to SLBs at higher densities than ICAM-1 (Fig. 1g), perhaps due to steric hindrance of 8 N-linked glycans on ICAM-1 compared to none on UCHT-1. When Ni–NTA percentage exceeded 9%, binding sites were no longer limiting, and UCHT-1 and ICAM-1 reached comparable densities (Fig. 1g and Extended Data Fig. 4b). Both proteins exhibited similar diffusion behavior on the SLBs, indicating free lateral mobility despite differences in molecular size (Extended Data Fig. 4c). At higher protein concentrations, densities reached a saturation plateau (Fig. 1g). Because fluorophore self-quenching can occur at high molecular densities to reduce emission intensity^30^, we tested whether this density saturation in UCHT-1 and ICAM-1 reflected binding saturation rather than quenching. By modulating the fraction of fluorescent lipids in the SLBs, we defined the linear, non-self-quenching range for each fluorophore (Fig. 1h,i). The saturation densities observed for UCHT-1 and ICAM-1 fell well within the linear range (Fig. 1j and Methods), confirming that QuEST-reported saturation arises from binding site occupancy rather than fluorophore quenching.

We next extended QuEST to quantify ligand dynamics on the SLBs related to IS formation. T cells were allowed to interact with SLBs containing UCHT-1 (36 molecules/µm^2^) and ICAM-1 (177 molecules/µm^2^) for 2, 10, or 20 min at 37 °C, followed by fixation and imaging at 24 °C. ICAM-1 rapidly accumulated into the peripheral supramolecular activation cluster (pSMAC) within the first 2 minutes, with a plateau reached by 10 and 20 min. In contrast, UCHT-1 continued to be delivered to the cSMAC, accumulating more slowly over time (Fig. 1k). These findings reveal distinct TCR dynamics with the high-affinity UCHT-1 compared to those triggered by lower-affinity pMHC, which peaked early and then decreased over time^1^. Live cell imaging at 37 ℃ under identical conditions yielded similar results (Extended Data Fig. 4d). Although single-tag intensities are lower at 37 °C (Extended Data Fig. 1d–f), quantification at 24 °C and 37 °C remained fully consistent after accounting for temperature effects (Extended Data Fig. 4e), reinforcing the robustness of QuEST when the tags are incorporated onto SLBs.

We next examined Jurkat cells, a widely used model for TCR signalosome biochemistry and visualization^31^. Using QuEST, we found that Jurkat cells achieved 70–90% of the final UCHT-1 and ICAM-1 densities observed in T cells; ICAM-1 accumulation kinetics were slower in Jurkat cells, consistent with lower levels of LFA-1^32^, whereas UCHT-1 kinetics were comparable (Fig. 1k and Extended Data Fig. 4f). These data suggest that Jurkat cells can approximate primary T cells for signalosome studies when given sufficient time to reach ICAM-1 saturation.

Quantitative measurements using TIRFM raise concerns that non-uniform illumination across the field of view may bias signal intensity. To address these concerns, we performed TIRF imaging across four fluorescence channels: 405 nm, 488 nm, 561 nm, and 647 nm. We prepared supported lipid bilayers (SLBs) containing 1% of each fluorescent DOPE variant (Atto 390, Atto 488, Atto 565, and Atto 647) and 96% DOPC. These SLBs were imaged using TIRFM under laser powers and camera exposure times comparable to those used in our experiments (Extended Data Fig. 5a). We quantified the relative mean fluorescence intensities (MFI) along both X and Y line profiles within 60 µm × 60 µm imaging regions. Across all channels, the intensity profiles exhibited bell-shaped distributions, with higher relative MFI at the center and lower values toward the edges (Extended Data Fig. 5b). The standard deviations (SD) for all channels were between 10% and 20%, indicating that the measured intensities can be approximated as the mean value with 10%–20% variance (Extended Data Fig. 5c). We further evaluated the variance with cell samples. T cells were cultured on SLBs containing UCHT-1 simultaneously labelled with AF405, AF488, AF568, and AF647 (36 molecules/µm^2^) and ICAM-1 (177 molecules/µm^2^) (Extended Data Fig. 5d). Identical regions were imaged at nine distinct positions across the field of view (Extended Data Fig. 5e,f). Relative to the central position, an SD of approximately 20% was defined for all channels (Extended Data Fig. 5g). These results demonstrate that illumination non-uniformity in our TIRFM system is limited and quantifiable, allowing its impact on quantitative analyses to be accounted for when we use the data in the imaging region.

### Calibration of TCR tags using engineered Jurkat cells

After the application of the quantification of ligands on the SLBs, we next sought to achieve an accurate estimation of the stoichiometry of the TCR signalosome using the QuEST. We first needed to determine the number of TCRs engaged in signaling in T cells. In principle, QuEST allows quantification of fluorescently tagged proteins, such as GFP, directly from their fluorescence, enabling a TCR–GFP strategy to measure total TCR abundance. In parallel, by staining TCR with UCHT-1 and quantifying the number of UCHT-1 bound per TCR, we can establish the quantitative relationship between UCHT-1 and TCR. This calibration then allows us to infer the number of non-GFP-tagged TCRs on the surface in the TIRF field in cells solely from UCHT-1 quantification afterward.

Because Jurkat cells are readily genetically modified and exhibit signaling dynamics comparable to T cells (Fig. 1k and Extended Data Fig. 4f), we used them to calibrate the quantitative relationship between TCR and UCHT-1. We generated Jurkat cells lacking endogenous TCR and expressing a 1G4 TCR–GFP fusion at a defined 1:1 GFP:TCR stoichiometry. In this background, CD8 was co-expressed, and CD4 was eliminated (Extended Data Fig. 6a). GFP-based quantification, however, requires careful correction: PFA fixation reduces GFP fluorescence (Extended Data Fig. 1i), and processing steps such as permeabilization and staining can lead to molecular loss. In addition, approximately 30% of intracellular GFP molecules reside in dark states^33^. We further found that although PFA fixation reduces the fluorescence of GFP, it does not alter the fraction of GFP in the dark state, as comparable numbers of single GFP molecules were detected before and after PFA fixation (Extended Data Fig. 7). All of these factors were incorporated into our quantitative framework.

To quantify the TCRs on the scanning cells, we cultured 1G4 TCR–GFP Jurkat cells on ICAM-1-coated SLBs without activating TCR signaling. We quantified 1G4 TCR densities in live cells (Live) and in cells subjected to fixation, permeabilization, and staining (FPS) (Fig. 2a and Extended Data Fig. 8a). To ensure that UCHT-1 bound exclusively to TCR and not to the SLBs, we used imidazole to block potential UCHT-1–SLB interactions during the staining phase (Supplementary Fig. 1). In the scanning Jurkat cells, we observed a ∼30% loss of TCRs during processing, and the measured UCHT-1:TCR ratio was 0.344 (Fig. 2b). Note that, while the UCHT-1 would have access to intracellular TCR, the use of TIRFM is expected to bias detection toward surface TCR in the IS.

**Fig. 2.**
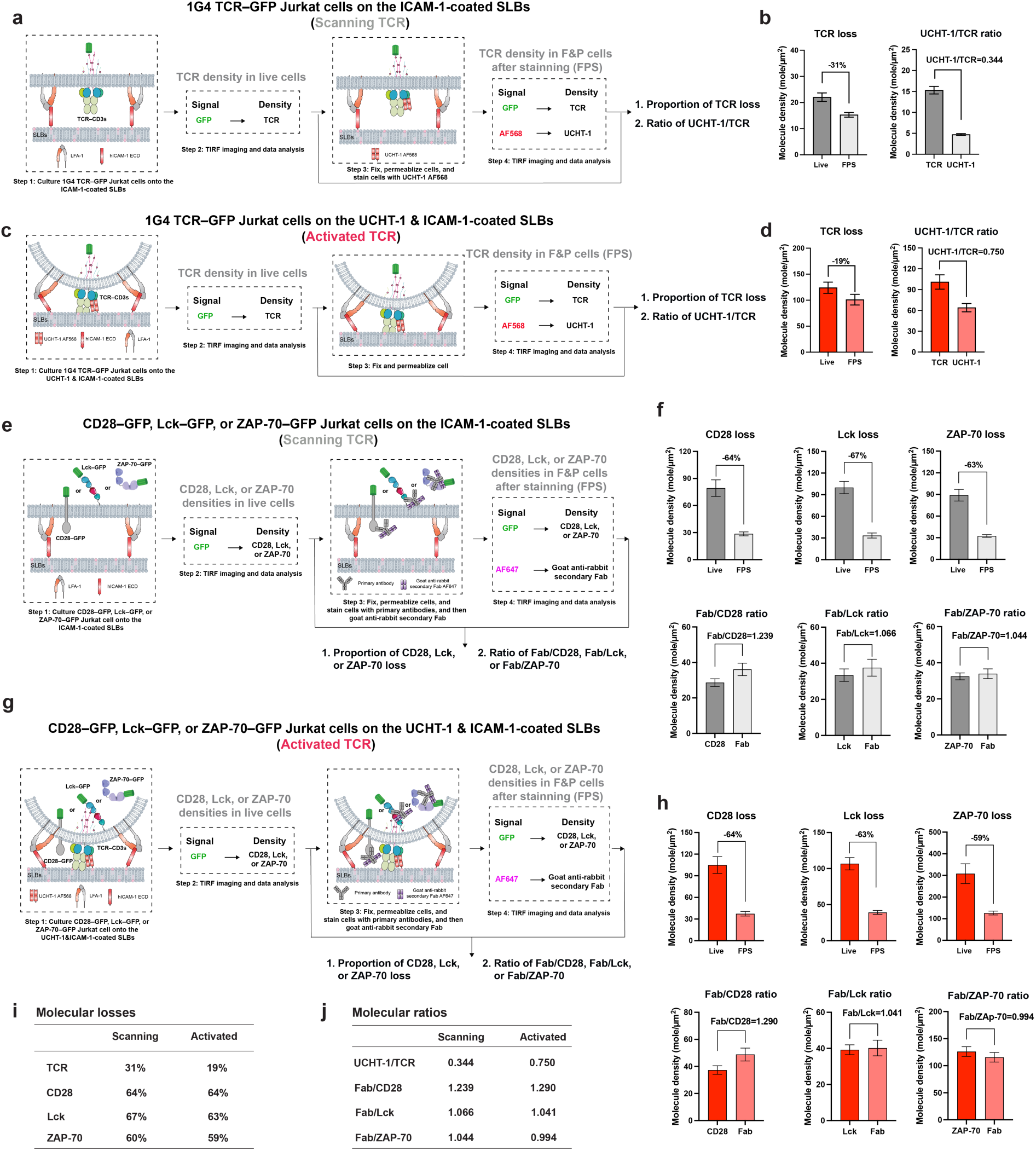
Quantification of molecular loss and molecular ratios for calibration of QuEST. **a**–**d**, During TCR scanning (**a**,**b**) or upon TCR activation (**c**,**d**), workflow (**a,c**) and quantification results (**b,d**) of TCR loss following cellular treatments and determination of the UCHT-1 to TCR ratio. **e**,**f**, During TCR scanning, workflow (**e**) and quantification results (**f**) of CD28, Lck, or ZAP-70 loss following cellular treatments and determination of the secondary Fab to CD28, Lck, or ZAP-70 ratios. **g**,**h**, Upon TCR activation, workflow (**g**) and quantification results (**h**) of CD28, Lck, or ZAP-70 loss following cellular treatments and determination of the secondary Fab to CD28, Lck, or ZAP-70 ratios. **i** and **j**, Quantified molecular loss (**i**) and corresponding molecular ratios (**j**) used to calibrate QuEST. Data in **b**,**d**,**f**, and **h** are representative of 2 replicates and presented as mean ± SEM from ≥30 individual cells.

We then quantified activated TCRs on cells activated by SLBs coated with UCHT-1 & ICAM-1 (Fig. 2c and Extended Data Fig. 8b). In activated Jurkat cells, TCR loss during processing decreased to ∼20%, likely reflecting increased receptor stability within the assembled signalosome. The UCHT-1:TCR ratio increased to 0.75 (Fig. 2d). Because receptor–ligand or epitope–antibody engagement is rarely complete, particularly in crowded environments such as the signalosome, ratios below unity are expected and consistent with known limitations of ligand accessibility. This did not affect our calibration, as we determined the number of TCRs by GFP quantification and subsequently derived the UCHT-1:TCR ratio.

With this TCR calibration in hand (Fig. 2b,d,i,j), we can now combine measurements of UCHT-1 surface density, obtained either from scanning cells on ICAM-1 only SLBs or from activated cells on ICAM-1 & UCHT-1 SLBs, with the experimentally determined UCHT-1:TCR ratio and TCR loss factor to infer the surface density of non-GFP tagged TCRs in T cells based on the UCHT-1 on the SLBs under scanning or activating conditions.

### Calibration of tags in TCR signalosome using engineered Jurkat cells

To determine the stoichiometry of signaling components within the TCR signalosome, we next quantified the numbers of key signaling molecules surrounding the TCR. As representative transmembrane, membrane-associated, and cytosolic signaling proteins, we selected CD28, Lck, and ZAP-70 for calibration. We generated Jurkat cell lines lacking the endogenous molecules and expressing single GFP-tagged CD28, Lck, or ZAP-70 (Extended Data Fig. 6c–d). Using the same experimental conditions employed to modulate TCR activation, we stained these cells with unlabelled primary antibodies followed by AF647-labelled goat anti-rabbit secondary Fab fragments (together the tag for the protein of interest) (Fig. 2e,g, and Extended Data Fig. 8c,d), enabling us to quantify molecular losses during FPS and to establish the quantitative relationship between secondary Fab tag signal and the original GFP signal in Jurkat cells. In scanning Jurkat cells, ∼60% of CD28, Lck, and ZAP-70 molecules were lost during FPS, and the measured Fab:molecule ratios were ∼1 (Fig. 2f,i,j). Upon TCR activation of Jurkats, molecular loss remained ∼60% for these molecules, and the Fab:molecule ratios similarly remained ∼1 (Fig. 2h,i,j). This ratio reflects several factors combined: primary antibody binding efficiency, secondary Fab binding efficiency, and the presence of multiple secondary Fab binding sites on each primary antibody. The greater loss of CD28, Lck, and ZAP-70 relative to TCR likely reflects their lower structural stability, as the TCR complex contains six CD3 chains and recruits numerous interaction partners that stabilize it within the signalosome. Importantly, we observed substantial similarity in the relevant calibration parameters across all molecule types except TCR (Fig. 2i,j). This motivated us to apply these parameters to five additional signaling components—CD8, CD45, PD-1, LAT, and PLCγ1.

With these molecular calibrations established, we can now quantify the densities of tag-free signaling proteins in scanning or activated primary T cells directly from the measured densities of secondary Fab, together with the experimentally determined Fab:molecule ratios and molecular loss factors to recover the original molecular densities and stoichiometries using identical conditions.

Applying single-molecule calibration data acquired at the glass interface to signals originating several nanometers inside the cell (e.g., GFP, ZAP-70, Lck, and PLCγ1) raises an additional concern: whether TIRFM illumination remains sufficiently uniform across the greater axial depth of the membrane-proximal signalosome, which may extend up to ∼30 nm from the surface. For GFP measurements, ZAP-70–GFP–expressing Jurkat cells were cultured on SLBs coated with UCHT-1 AF488 (36 molecules/µm^2^) and ICAM-1 (177 molecules/µm^2^). Cells were stained with an anti-ZAP-70 primary antibody followed by an AF647-labelled secondary Fab fragment. We quantified the density of clustered Fabs and, using the calibrated Fab:ZAP-70 molecular ratio (Fig. 2j), determined the density of clustered GFP. This enabled quantification of the fluorescence intensity of single GFP molecules within the TCR signalosome (Extended Data Fig. 9a). For the quantification of the fluorescence intensity of purified GFP protein adsorbed onto the coverslip (Extended Data Fig. 9b), and to obtain a comparable spatial pattern used in cells, here we used B(c)–GFP, in which a monomeric GFP is fused to a tandem GmhA scaffold^28^. We confirmed that GFP fluorescence within the TCR signalosome is comparable to that measured on the coverslip (Extended Data Fig. 9c). We next compared the mean fluorescence intensities of single Fab in TCR signalosomes (by staining ZAP-70 and diluting the AF647-labelled Fab by 2000-fold) (Extended Data Fig. 10a) with single secondary Fab molecules adsorbed to the coverslip directly (Extended Data Fig. 10b). The mean fluorescence intensities of single Fab molecules in the TCR signalosome and at the coverslip were indistinguishable (Extended Data Fig. 10c). These data demonstrate that TIRFM provides sufficiently uniform fluorescence excitation and detection across an axial range of at least 0–30 nm, supporting accurate quantification of membrane-proximal intracellular molecules. Insensitivity to depth in this range is expected in practice^34^.

We confirmed that QuEST yields relative errors below 5% when quantifying TCR and signaling molecule densities in Jurkat cells (Extended Data Fig. 11).

### Mapping the dynamic stoichiometries of the TCR signalosome

With all calibration parameters for TCR and the eight other signaling molecules in place, we were poised to map the stoichiometries of the TCR signalosome in human primary T cells, which were activated in vitro and maintained with IL-2 for 10 days, during scanning for antigen and at different stages following TCR engagement, with high confidence in minimizing error. We first investigated T cells scanning SLBs containing only ICAM-1 (177 molecules/μm^2^) (Fig. 3a). Under these conditions, an average of ∼43 TCRs/µm^2^ and the signaling molecules of interest (bulk) were broadly distributed across the T cell–SLBs interface (Fig. 3b and Supplementary Fig. 2a). We measured the molecular (X) densities across the entire scanning interface and calculated X:TCR ratios of 5.4 CD8, 0.9 CD28, 10.2 CD45, 0.1 PD-1, 3.4 Lck, 1.2 ZAP-70, 4.2 LAT, and 0.6 PLCγ1 per TCR (Fig. 3c). Due to the inability to laterally resolve signalosomes in the scanning interface, the measurements provide the lateral density of components to a depth of ∼100 nm with no implications for assembly. A recent study reported that certain molecules are pre-assembled into TCR microclusters prior to antigen engagement^35^. Accordingly, we defined the brightest region of TCR fluorescence as the pre-cluster during the TCR scanning state (Supplementary Fig. 2b). Within these regions, we measured an average density of ∼99 TCRs/µm². The corresponding molecule:TCR ratios were 4.1 for CD8, 0.4 for CD28, 6.4 for CD45, 0.06 for PD-1, 2.7 for Lck, 0.9 for ZAP-70, 2.8 for LAT, and 0.6 for PLCγ1 (Fig. 3d).

**Fig. 3.**
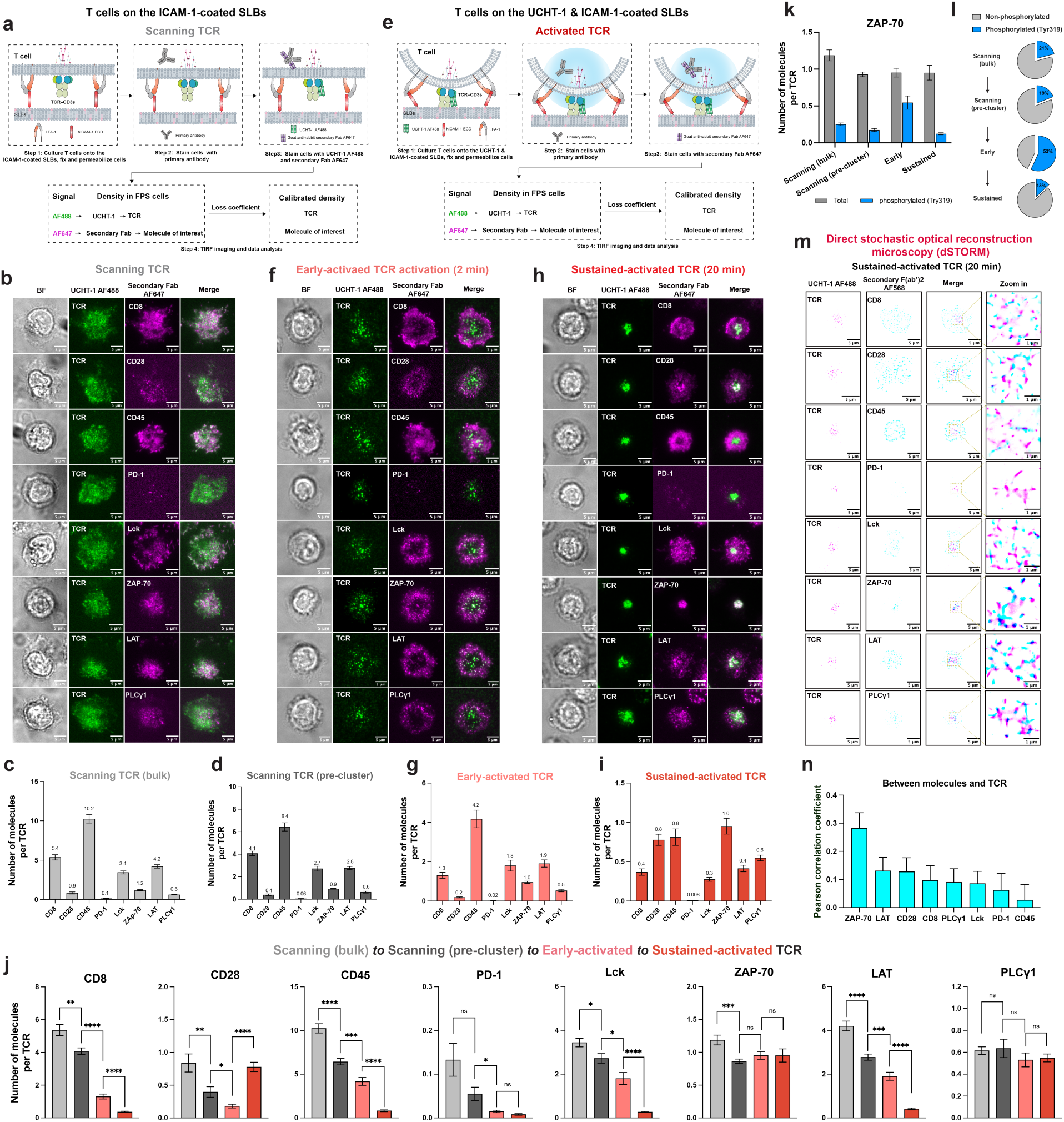
Mapping the spatiotemporal stoichiometry of the TCR signalosome. **a**–**d**, Workflow (**a**), representative TIRF images (**b**), and quantified bulk (**c**) and pre-cluster (**d**) results of the number of signaling molecules per TCR during the scanning states. **e**–**i**, Workflow (**e**), representative TIRF images at 2 min (**f**) and 20 min (**h**), and quantified results at 2 min (**g**) and 20 min (**i**) of the number of signaling molecules per TCR upon TCR activation. **j**, Dynamic stoichiometry of the TCR signalosome across different activation states. **k**,**l**, Number of phosphorylated and total (pan) ZAP-70 molecules per TCR (**k**) and proportions of phosphorylated and non-phosphorylated ZAP-70 (**l**) at different states. **m**, Representative dSTORM images of the cSMAC during sustained TCR activation. **n**, Pearson correlation coefficients between signaling molecules and TCR based on dSTORM imaging. Data are presented as mean ± SEM. Statistical significance was assessed using an unpaired two-tailed Student’s *t*-test (\*\*\*\**P* ≤ 0.0001; \*\*\**P* ≤ 0.001; \*\**P* ≤ 0.01; \**P* ≤ 0.05; ns, not significant). For each condition, ≥30 individual cells were analyzed, except for the data for scanning (pre-cluster), where ≥30 signaling regions from at least five individual cells were analyzed, and in panel **m**, where ≥20 signaling regions from at least three individual cells were analyzed. Data are representative of three replicates using T cells from three blood donors.

The TCR density in the pre-cluster region was 2.3-fold higher than that in the bulk region. Thus, if the molecule:TCR ratio in the bulk region is substantially lower than 2.3 times the ratio in the pre-cluster region, this suggests preferential pre-assembly of that molecule within TCR pre-clusters prior to activation. We inferred that PLCγ1, Lck, CD8, and ZAP-70 exhibit significant pre-assembly, whereas LAT, CD45, PD-1, and CD28 do not.

We next activated T cells by culturing them on the SLBs containing UCHT-1 AF488 (36 molecules/μm^2^) and ICAM-1 (177 molecules/μm^2^), enabling the formation of immunological synapses (Fig. 3e). To capture the molecular dynamics during activation, we mapped stoichiometries at both early (2 min) and sustained (20 min) activation stages. At the early activation stage, consistent with previous reports, TCRs accumulated in microclusters (Fig. 3f and Supplementary Fig. 2c). TCR microclusters averaged ∼208 TCRs/µm^2^ with X:TCR ratios of: 1.3 CD8, 0.2 CD28, 4.2 CD45, 0.02 PD-1, 1.8 Lck, 1.0 ZAP-70, 1.9 LAT, and 0.5 PLCγ1 per TCR (Fig. 3g). By 20 min, most of the TCR and ZAP-70 concentrated in the single cSMAC where other signaling molecules displayed moderate concentration in the cSMAC (CD8, CD28, PD-1, Lck, LAT, and PLCγ1) or exclusion from the cSMAC (CD45) (Fig. 3h and Supplementary Fig. 2d). The cSMACs contained an average of ∼811 TCRs/µm^2^ with stoichiometries of 0.4 CD8, 0.8 CD28, 0.8 CD45, 0.008 PD-1, 0.3 Lck, 1.0 ZAP-70, 0.4 LAT, and 0.6 PLCγ1 per TCR (Fig. 3i). To better characterize the molecular dynamics underlying T cell activation, we further quantified the molecular densities within and outside the microclusters (Extended Data Fig. 12a,c) as well as the cSMAC (Extended Data Fig. 12b,c) during T cell activation. These measurements provide direct evidence for the selective recruitment or exclusion of molecules at sites of TCR signaling. Altogether, we presented the dynamics in stoichiometries across the T cell activation, showing the adaptability of the TCR signalosome machinery when facing various stimulations (Fig. 3j).

Expected and unexpected features emerged from the analysis of the dynamic TCR signalosome stoichiometries during T cell activation. First, Lck is rapidly recruited to microclusters upon initial TCR activation. Subsequently, its density increases within the cSMAC. However, comparable levels of Lck are also observed outside the cSMAC, suggesting that at later stages of TCR activation, a high stoichiometry of Lck is not essential for sustaining TCR signaling (Fig. 3i,j, and Extended Data Fig. 12c). Second, CD28 behavior in the absence of any ligand on the SLBs was biphasic with random distribution during scanning and not in the TCR pre-clusters, formation of positionally distinct microclusters from TCR at 2 min, and significant co-localization with TCR in the cSMAC at 20 min (Fig. 3b,f,h, and Extended Data Fig. 12c). Clustering of CD28 without a ligand on the SLBs likely results from cis interaction with CD80/CD86 in tubular invaginations, as recently described^36^. Third, as expected, CD45 is reduced in the cSMAC with ∼1076 molecules/μm^2^ outside and ∼631 molecules/µm^2^ inside (Fig. 3h and Extended Data Fig. 12c). Surprisingly, there are still ∼0.8 CD45 per TCR within the cSMAC exceeding all other signaling molecules except ZAP-70 (Fig. 3i). Fourth, as expected, ZAP-70 dominates in both abundance and stoichiometry with TCR compared with other signaling molecules. Surprisingly, although the TCR complex contains 10 ITAMs that may recruit ZAP-70, only 1 ZAP-70 per TCR is detected in both the early TCR microclusters and in the sustained cSMAC (Fig. 3g,i,j, and Extended Data Fig. 12c). This may reflect the high density of CD45 in the cSMAC, which can accelerate ZAP-70 dissociation from ITAMs^37^. The stoichiometric ratio of ZAP-70 to TCR remains approximately one across scanning (pre-cluster), early-triggering, and sustained states, such that it operates similarly to receptor kinases that typically have one kinase domain per receptor. However, we have no evidence that the ZAP-70 is bound to the TCR during scanning (bulk), and the relatively high ratio during this phase suggests that sufficient ZAP-70 to support early microcluster formation is within less than 100 nm prior to any TCR engagement. Lastly, the close correspondence between CD8 and Lck stoichiometries in the microclusters and cSMAC (Fig. 3g,i) is consistent with a substantial fraction of Lck being associated with CD8, as expected^38^.

To further refine the dynamic stoichiometries, we incorporated post-translational modifications into the QuEST by quantifying phosphorylated signaling molecules during TCR activation. We focused on ZAP-70, given its central role as discussed above (Fig. 3f–j). Intriguingly, before activation, ZAP-70 existed in a basal phosphorylated pool of ∼20% at scanning bulk state. Although ZAP-70 is pre-assembled to TCR pre-clusters during TCR scanning, its phosphorylated level is not significantly altered (Fig. 3k,l).

During the early triggering stage, the fraction of phosphorylated ZAP-70 increased sharply. However, during sustained activation, although the number of phosphorylated ZAP-70 molecules in the cSMAC increased, the phosphorylation percentage reverted to below basal levels (Fig. 3k,l), indicating that ZAP-70 phosphorylation peaks rapidly during early activation, as expected from bulk analysis of T cell responses to TCR crosslinking^39^. These data suggest that ZAP-70 functions as a sensitive gatekeeper of TCR activation by modulating its phosphorylation levels.

To visualize the nanoarchitecture underlying these diffraction-limited measurements, we complemented QuEST-based quantification with super-resolution imaging using direct stochastic optical reconstruction microscopy (dSTORM)^17^. Although dSTORM is not suitable for molecular quantification due to the complex behavior of individual fluorophores, it offers ∼20-nm spatial resolution and may shed light on the actual footprint of molecules in the cSMAC, which we suspect is a mosaic of nanoscale clusters. We found that in the cSMAC during sustained activation, ZAP-70 most strongly colocalized with TCR, followed by LAT, CD28, CD8, PLCγ1, Lck, PD-1, and CD45 (Fig. 3m,n). Despite the presence of abundant CD45 molecules in the cSMAC (Extended Data Fig. 12c), the dSTORM data suggest that they stay far from TCR on the nanoscale. Notably, Lck localized at the periphery rather than the center of the signalosome (Fig. 3m), suggesting that during sustained activation—when phosphatases such as CD45 are spatially excluded from the signalosome core—the best location for Lck may be at the interface between the CD45 (needed to remove inhibitory C-terminal phosphorylation of Lck) and TCR (Fig. 3n) to phosphorylate ITAMs.

By integrating dynamic measurements and quantification, we estimated the spatiotemporal stoichiometry of the TCR signalosome to quantitatively elucidate the stoichiometric ratios, spatial organization, and post-translational modifications of signaling molecules during TCR activation (Extended Data Fig. 13). Finally, we confirmed that the relative errors in QuEST-based quantification of molecule numbers on both the cell surface and SLBs across activation stages remained below 5% (Extended Data Fig. 14).

In addition, FRET is also a possible source of quenching. We avoided FRET by using dye combinations such as AF488 and AF647 in most of the QuEST applications, where there is no overlap of AF488 emission with AF647 excitation, precluding FRET (See Methods). To rule out artefacts from multistep antibody staining, we repeated experiments using fluorescent primary antibodies instead of the nonfluorescent-primary/fluorescent-secondary Fab scheme and observed highly consistent imaging patterns (Supplementary Fig. 3a,b). To ensure high-fidelity quantification using QuEST, we verified that without primary antibodies, the secondary Fab exhibited no cross-reactivity toward signaling molecules or UCHT-1 on the SLBs (Supplementary Fig. 4a,b).

The unexpected result that CD28 clustering in the cSMAC in the absence of a trans CD28 ligand led us to further investigate whether PD-1 engagement by PD-L1 would impact this process, given that PD-L1 can interact with CD80^40^.

### PD-1 engagement antagonizes CD28 clustering in the cSMAC

Since its discovery^41^, the inhibitory receptor PD-1 and its ligand-induced suppression of TCR signaling have been extensively investigated^42,43^, revealing substantial therapeutic potential in cancer immunotherapy^44^. Here, we quantified how PD-1 engagement by PD-L1 reshapes the stoichiometric architecture of the TCR signalosome. Based on previous studies on APC model cell lines and physiologically relevant PD-L1 densities used in reconstitution assays, PD-L1 surface densities on activated APCs are estimated to lie at a high level (∼10^2^–10^3^ molecules/µm^2^)^36,45^. We quantified PD-L1 (extracellular domain throughout the study; sequence in Methods) on the SLBs (Extended Data Fig. 15). T cells were introduced onto the SLBs functionalized with UCHT-1 AF488 (36 molecules/µm^2^), ICAM-1 (177 molecules/µm^2^), and PD-L1 (400 molecules/µm^2^) (Fig. 4a), enabling determination of the stoichiometry of the TCR signalosome by PD-1 engagement (Fig. 4b–d). PD-1 engagement led to ∼702 TCRs/µm^2^ in the cSMAC with a stoichiometry of 0.9 CD8, 0.08 CD28, 0.4 CD45, 0.2 PD-1, 0.3 Lck, 1.4 ZAP-70, 1.5 LAT, and 0.3 PLCγ1 per TCR (Fig. 4c). Single-cell analyses frequently define significant responses based on relative thresholds (e.g., ∼60% of baseline), further illustrating that percentage changes in signaling molecule activity are interpreted as biologically meaningful^46^. Using a 60% relative change threshold here, we found that PD-1 engagement led to marked increases in PD-1 (+2400%), LAT (+275%), and CD8 (+125%), while CD28 decreased by 90% in the TCR signalosome (Fig. 4e). The marked reduction of CD28 within the cSMAC supports previous findings that CD28 is a primary target of PD-1-mediated inhibition^47^, but this is the first data we are aware of that supports the hypothesis that PD-1 ligation inhibits CD28 signaling mediated by cis interactions with CD80/CD86 expressed on the T cell^36^. By contrast, CD45, Lck, ZAP-70, and PLCγ1 were less affected (Fig. 4d,e), indicating molecule-specific sensitivities to PD-1 inhibitory signaling (Fig. 4e).

**Fig. 4.**
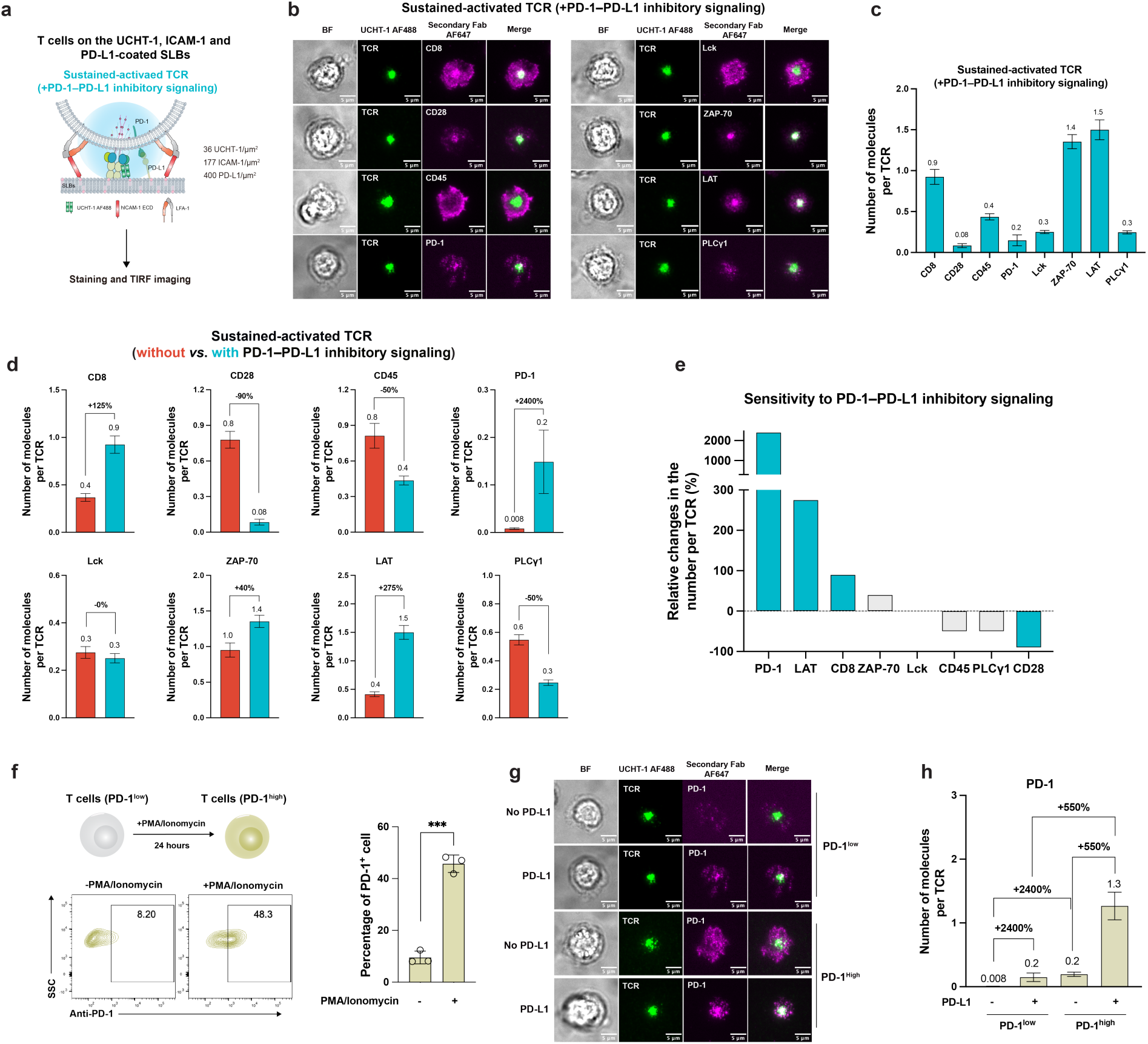
PD-1–PD-L1 inhibitory signaling reshapes the stoichiometry of the TCR signalosome by antagonizing CD28 clustering in the cSMAC. **a**–**c**, Workflow (**a**), representative TIRF images (**b**), and quantification results (**c**) of the number of signaling molecules per TCR following introduction of PD-1–PD-L1 inhibitory signaling in T cells. **d**, Stoichiometry of the TCR signalosome under PD-1–PD-L1 inhibitory signaling. **e**, Quantified sensitivity of signaling molecules within the TCR signalosome to PD-1–PD-L1 inhibitory signaling. **f**, Generation of T cells expressing low or high levels of PD-1. **g**,**h**, Representative TIRF images (**g**) and corresponding quantification results (**h**) of the number of signaling molecules per TCR in T cells expressing low or high levels of PD-1, with or without PD-1–PD-L1 inhibitory signaling. Data in **f** are presented as mean ± SD; data in **c**,**d**, and **h** are presented as mean ± SEM. In **c**,**d**, and **h**, ≥30 individual cells were analyzed per condition. Statistical significance was assessed using an unpaired two-tailed Student’s *t*-test (\*\*\**P* ≤ 0.001). Data in **c**,**d**, and **h** are representative of two replicates using T cells from two blood donors. Data in **f** are representative of three replicates using T cells from three blood donors.

Interestingly, although PD-1 ligand engagement substantially increased PD-1 recruitment, the PD-1:TCR stoichiometric ratio remained low (∼0.2). Meanwhile, ZAP-70 and LAT continued to dominate over PD-1 in ratios. These observations suggest that the maximal inhibitory potential of PD-1 may not be realized in these T cells, which express relatively low endogenous PD-1, consistent with PD-1 being inducible rather than constitutively high in effector T cells. In physiological contexts such as tumors or chronic infection, PD-1 expression can be markedly upregulated^48^. To mimic these conditions, we increased PD-1 expression on T cells (Fig. 4f) and repeated the PD-L1 stimulation experiments with PD-1^high^ cells. In PD-1^high^ T cells, PD-1 recruitment to the signalosome increased dramatically (Fig. 4g). Without PD-L1, PD-1 recruitment rose by 2400% relative to PD-1^low^ cells; with PD-L1, the increase was 550%, yielding 1.3 PD-1 molecules per TCR in the signalosome (Fig. 4h). These results reveal the strong dependence of PD-1 inhibitory stoichiometry on receptor expression levels and the presence of its ligand.

### CD28 engagement by CD86 in trans promotes TCR proximal signals

CD28 ligation by CD86 or CD80 in trans is well known to enhance TCR activation and plays a central role in strengthening T cell-mediated anti-tumor immunity^49^. How this relates to the CD28 clustering driven by intrinsic mechanisms like cis interactions with CD86 is not known. CD86 is a highly expressed ligand on mature dendritic cells with a molecular density over 100 molecules/µm^2^ quantified previously^50^, we therefore established SLBs presenting CD86 (extracellular domain throughout the study; sequence in Methods) at 400 molecules/µm^2^ (Extended Data Fig. 16). T cells were then introduced onto the SLBs functionalized with UCHT-1 AF488 (36 molecules/µm^2^), ICAM-1 (177 molecules/µm^2^), and CD86 (400 molecules/µm^2^) (Fig. 5a), enabling determination of the TCR signalosome stoichiometry in the presence of CD28 engagement in trans (Fig. 5b–d). CD28 engagement led to ∼486 TCRs/µm^2^ in the cSMACs with stoichiometries of 1.1 CD8, 1.3 CD28, 0.5 CD45, 0.01 PD-1, 0.4 Lck, 1.4 ZAP-70, 1.8 LAT, and 0.4 PLCγ1 per TCR (Fig. 5c). Applying a 60% relative-change threshold, CD28 costimulation induced pronounced increases in LAT (+350%), CD8 (+175%), and CD28 (+63%) in the TCR signalosome, which are assumed to promote the TCR proximal signaling. In contrast, CD45, PD-1, Lck, ZAP-70, and PLCγ1 were minimally affected (Fig. 5d), reflecting molecule-specific responsiveness to CD28 signaling engagement (Fig. 5e). Upon ligand engagement, the increase in CD28 abundance within the TCR signalosome is less pronounced than that of PD-1, likely due to the competition between the ligands on the SLBs with CD86 expressed on T cell surface^36^ and/or the competition between CD28 with CTLA-4 expressed on the T cell surface^51^. It is striking that PD-1 and CD28 engagement had similar effects on LAT and CD8 recruitment, suggesting potential shared agonist activity of PD-1 or CD28 engagement on TCR signaling in addition to the disruption of CD28 signaling by PD-1 engagement.

**Fig. 5.**
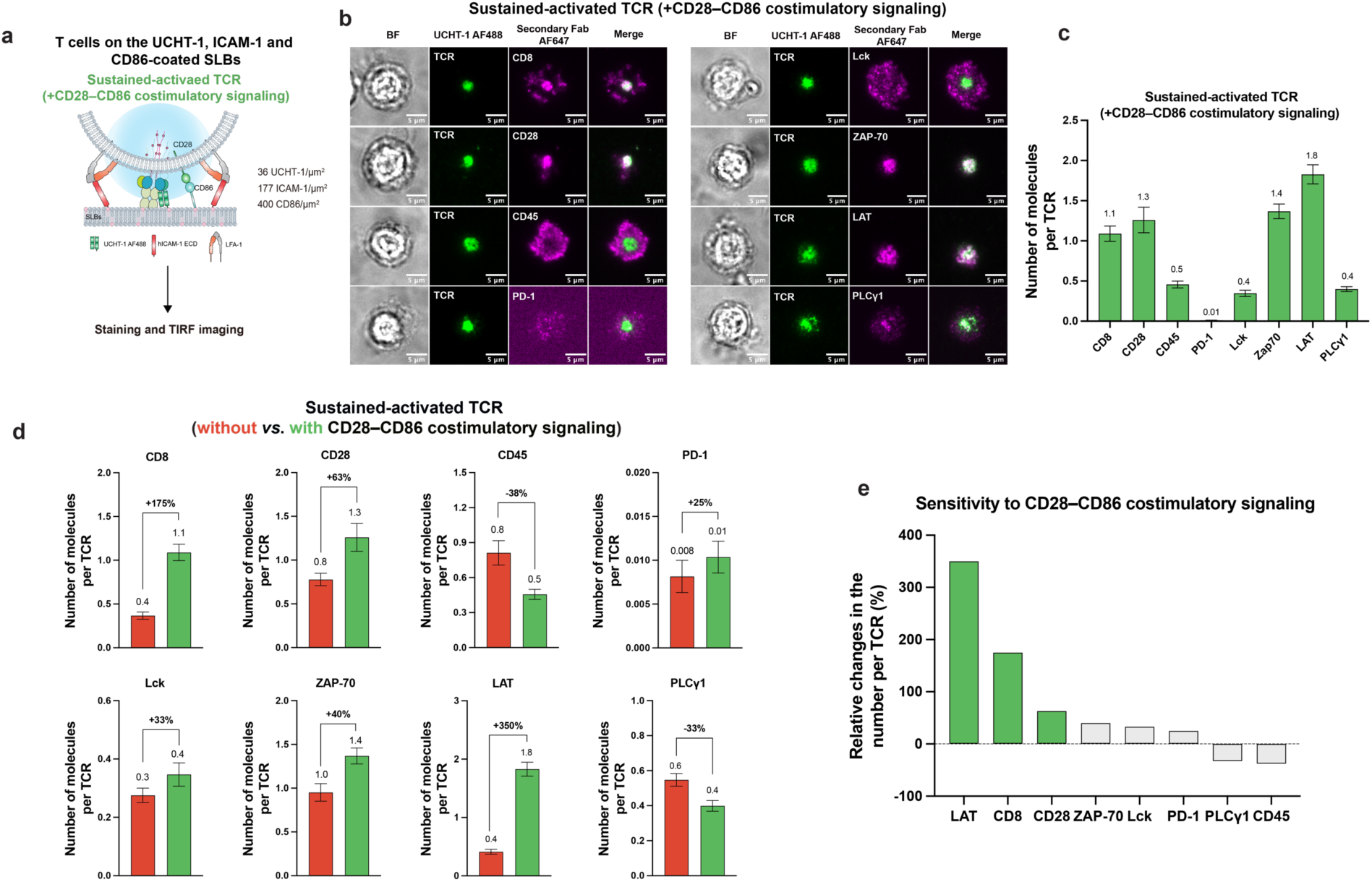
CD28–CD86 costimulatory signaling reshapes the stoichiometry of the TCR signalosome by promoting TCR proximal signals. **a**–**c**, Workflow (**a**), representative TIRF images (**b**), and quantification results (**c**) of the number of signaling molecules per TCR following introduction of CD28–CD86 costimulatory signaling in T cells. **d**, Stoichiometry of the TCR signalosome under CD28–CD86 costimulatory signaling. **e**, Quantified sensitivity of signaling molecules within the TCR signalosome to CD28–CD86 costimulatory signaling. Data in **c** and **d** are presented as mean ± SEM. In **c** and **d**, ≥30 individual cells were analyzed per condition. Data in **c** and **d** are representative of two replicates using T cells from two blood donors.

## Discussion

In this study, we developed the QuEST method, which overcomes long-standing challenges in direct, accurate estimation of molecular numbers and stoichiometric ratios within dense and crowded signaling complexes detected by immunofluorescence TIRFM. Through careful calibration and minimizing measurement errors from multiple sources, QuEST enabled the accurate extrapolation of the spatiotemporal stoichiometry of the TCR signalosome across scanning, early-triggered, and sustained activation, coinhibited, and costimulated states (Fig. 6).

**Fig. 6.**
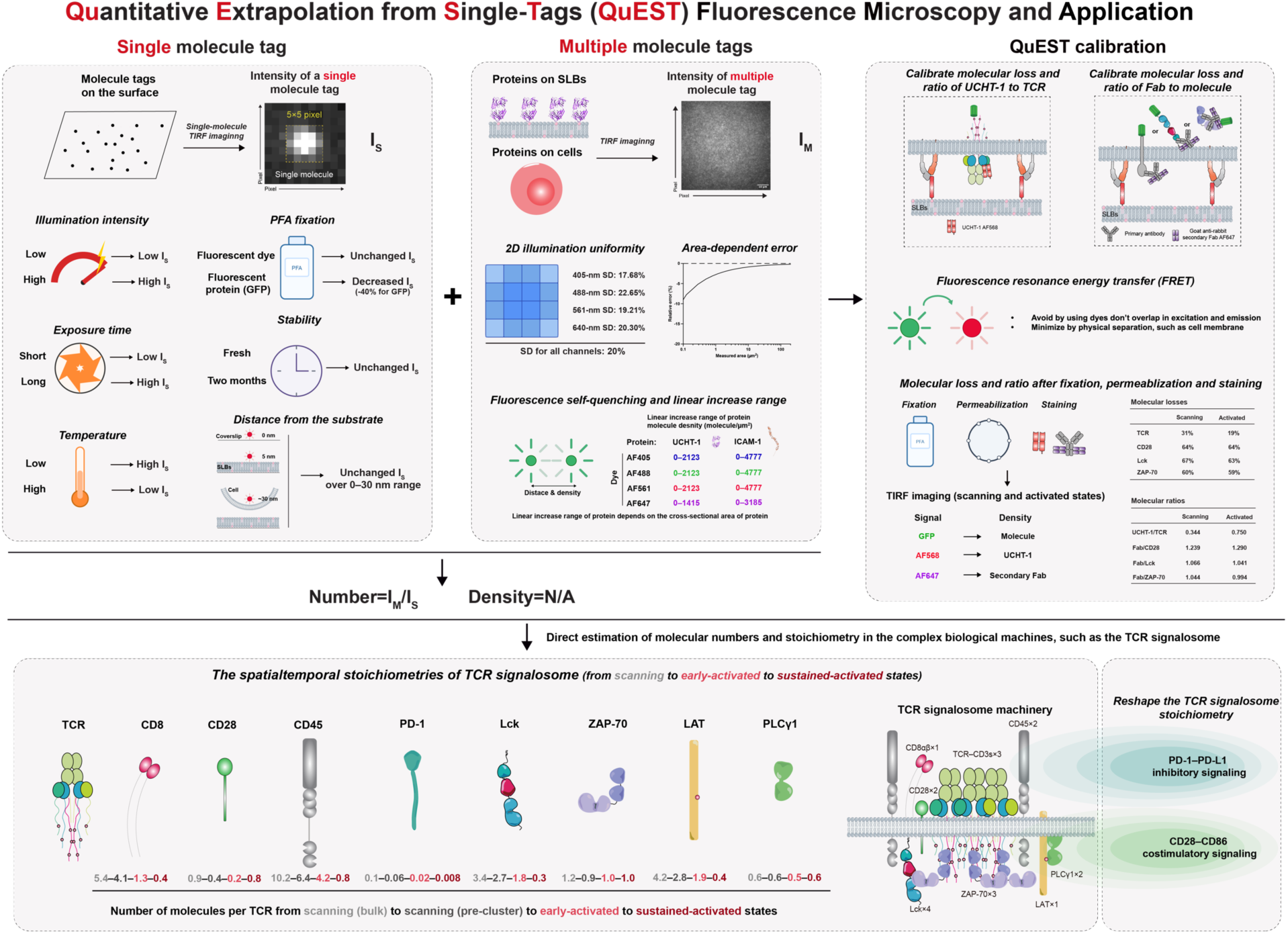
Overview of the QuEST method and its application in direct estimation of stoichiometries of TCR signalosome machinery.

TCR signalosome stoichiometry reveals quantitative principles that govern TCR signaling. The stoichiometric composition of the signalosome undergoes substantial reorganizations upon activation. ZAP-70 and LAT consistently dominate intracellular signaling components. In particular, the stoichiometric ratio of ZAP-70 to TCR remains approximately 1:1 across scanning, early-triggered, and sustained states. This one-to-one coupling of ZAP-70 and TCR over time suggests a unique mechanism in which ZAP-70 is closely aligned with changes in both the quantity and density of TCR, while modulating the phosphorylation of ZAP-70 to set the TCR triggering threshold. CD28 and PD-1 each reshape the signalosome in distinct and common ways: CD28 and PD-1 engagement both markedly enrich LAT and CD8 in the cSMAC, while reducing CD45, thereby shifting the kinase–phosphatase balance toward activation of TCR proximal signals. PD-L1 on the SLBs increased PD-1 clustering in the cSMAC and suppressed the T cell autonomous CD28 clustering. T cells expressing higher levels of PD-1, consistent with chronically stimulated T cells, could reach similar levels of PD-1 to TCR and ZAP-70. These quantitative dependencies offer mechanistic explanations for context-dependent responses to costimulation and checkpoint inhibition.

Our results provide a quantitative foundation for modelling TCR triggering, demonstrating that signaling output is governed not only by molecular presence but by precise stoichiometric ratios dictating competition and cooperation within the nanoscale signalosome. These findings have direct implications for T cell engineering. QuEST offers a platform to evaluate how engineered TCRs, CARs, or signaling motifs reconfigure the signalosome, enabling rational tuning of LAT or ZAP-70 enrichment, CD45 exclusion, or PD-1 accessibility. The stoichiometric basis of PD-1 inhibition further highlights why PD-1 blockade is most effective in PD-1^high^ states and suggests new approaches for checkpoint modulation.

Furthermore, extending the QuEST to dissect other T cell subsets (e.g., αβ versus γδ T cells, CD8⁺ versus CD4⁺ T cells) and to immune cells with unique signaling architectures (e.g., B cells, NK cells, macrophages, dendritic cells) will further illuminate how stoichiometric signaling principles vary across the immune system. Nonetheless, QuEST establishes a robust quantitative pipeline for dissecting nanoscale signaling networks and will also be applicable to the quantification of fluorescence signals that can be acquired by TIRFM. By illuminating the molecular architecture underlying TCR activation, inhibition, and costimulation, this work provides a framework for precision-guided T cell immunotherapy, including TCR-T optimization and improved strategies for PD-1 blockade.

Finally, this study has some limitations. Although we quantified the numbers and densities of TCR and associated signaling molecules within the cSMAC, the stoichiometry was reconstructed from quantitative molecular ratios in the colocalized regions rather than by directly visualizing single molecules within intact signalosomes. Addressing this will require future methodologies with both higher effective resolution and in situ molecular counting. For example, qPAINT is a super-resolution method^26^ with the potential to count molecules within crowded signaling complexes that has been applied to T cells but not to TCR stoichiometric analysis^52,53^. While we have adopted an approach that extrapolates single-molecule fluorescence signals from conventional-resolution imaging data to robustly quantify components in crowded signalosomes directly from diffraction-limited images, our method does not require the long acquisition times or complex experimental reagents associated with qPAINT. The calibration framework we have developed could also be applied to improve qPAINT-based methods, for example, by enabling more accurate quantification of molecular loss and antibody-targeting efficiency. This is particularly important because substantial corrections are often required to relate tags applied to fixed and permeabilized samples to the true molecular density or target abundance present in live cells. We have solved this problem using an engineered cell line, Jurkat, in which GFP-tagged proteins can be quantified directly. TCR signalosomes can be used to calibrate more universal tags^54^ throughout the sample preparation process. Notably, the flat-top illumination strategy^55^ used in qPAINT imaging could also be implemented in our approach to reduce two-dimensional intensity variance, which we have quantified in our system. It is worth noting that certain regions of the cSMAC in CD8⁺ T cells correspond to extracellular vesicles, where signaling is terminated, and the TCR is removed from the T cell^56,57^. This is extensively documented as a rapid process in CD4^+^ T cells, whereas the timing of TCR ectocytosis in CD8^+^ T cells appears to be delayed, as the cSMAC is used to direct exocytosis into the IS prior to disengaging from the target through ectocytosis. This phenomenon may, to some extent, influence the estimation of the functional stoichiometries of the TCR signalosome. In addition, our TCR activation system relied on UCHT-1, a high-affinity CD3ε antibody, rather than pMHC ligands. Estimation of the stoichiometry of TCR triggered by distinct pMHCs, agonists, and antagonists would provide valuable insight into how T cells quantitatively discriminate self from non-self using the TCR signalosome machinery, a direction to be pursued in our future studies.

## Acknowledgements

We thank S. Valvo for support with the SLB platform, L. Chen for recombinant proteins, E. Kurz for lab management, and E. Jenkins for 1G4 TCR plasmid. We thank the staff, especially Y. Zhang and K. Korobchevskaya, in the Oxford-Zeiss Centre of Excellence for Biomedical Imaging. We thank Jonathan Webber, Manager of the Flow Cytometry Facility, for his assistance with flow cytometry troubleshooting and cell sorting. We thank W. White, C. J. Kim, and D. Baker for generously providing purified B(c)–GFP protein. We thank E. Smith, A. Armache, J. Waite, K. Olson, L. Haber, N. James, R. McKay, C. Meagher, D. Skokos, A. Ambegaonkar, and H. Tan at Regeneron Pharmaceuticals for suggestions on the manuscript. The TIRF microscope was purchased with funds from Wellcome PRF 100262Z/12/Z. We thank L. Gu at the Institute of Biophysics, Chinese Academy of Sciences, for helpful suggestions. We would also like to thank the anonymous blood donors who contributed to this study. We thank AlphaFold3 for its service in structure prediction. We appreciate BioRender for providing graphical elements used in figures. We thank Fiji for providing imaging analysis. This work was supported by a research agreement with Regeneron Pharmaceuticals, Wellcome Collaborative award (224343/Z/21/Z), the Kennedy Trust for Rheumatology Research, and the Chinese Academy of Medical Sciences (CAMS) Innovation Fund for Medical Science (CIFMS) (2024-I2M-2-001-1).

## Declaration of generative AI and AI-assisted technologies

During the preparation of this work, the authors used ChatGPT to improve the clarity and language of the manuscript. The authors reviewed and edited the output as necessary and take full responsibility for the content of the publication.

## Author contributions

P.F. conceived and designed the project with the guidance and supervision of M.L.D. P.F. performed experiments and data analyses, generated figures, and drafted the first manuscript. M.L.D. co-wrote the manuscript. P.F. and M.L.D. approved the final version of the manuscript.

## Methods

### Materials and Reagents

RPMI 1640 medium (Gibco, Cat#31870074), DMEM medium (ThermoFisher, Cat#11965092), Opti-MEM^TM^ medium (Gibco, Cat#11058021), Expi293^TM^ Expression Medium (ThermoFisher Scientific, Cat#A1435101), RosetteSep^TM^ human CD8^+^ T cell enrichment cocktail (Stemcell, Cat#15023), human T cell activating CD3/CD28 Dynabeads (ThermoFisher, Cat#11132D), recombinant human IL-2 (Peprotech, Cat#200-02), coverslip (Schott technical glass solutions GmbH, Cat#1472315), bottomless 6 channel slide (IBIDI, Cat#80608), Alexa Fluor 405 dye (AF405, ThermoFisher, Cat#A30000), Alexa Fluor 488 dye (AF488, ThermoFisher, Cat#A20000), Alexa Fluor 568 dye (AF568, ThermoFisher, Cat#A20003), Alexa Fluor 647 dye (AF647, ThermoFisher, Cat#A20006), Atto 390 DOPE (ATTO-TEC, Cat#AD 390-161), Atto 488 DOPE (ATTO-TEC, Cat#AD 488-165), Atto 565 DOPE (ATTO-TEC, Cat#AD 565-161), Atto 647 DOPE (ATTO-TEC, Cat#AD 647-161), DOPC lipid (Avanti Polar

Lipids, Cat#850375), Ni–NTA lipid (Avanti Polar Lipids, Cat#790404), viability dye (Invitrogen, Cat#65-0865-14), BD fixation/permeabilization kit (BD, Cat#554714), cysteamine (Merck, Cat#A30070), glucose oxidase (Merck, Cat#A49180-250), polyethylenimine (Polysciences, Cat#49553-93-7), PCR kits (New England Biolabs, Cat#M0494S), infusion kit (Takara, Cat#638947), plasmid miniprep kit (New England Biolabs, Cat#T1110L), DNA gel extraction kit (New England Biolabs, Cat#T1120L), spin desalting column (Thermo Scientific, Cat#89877), grRNA for CRISPR Cas9 (Integrated DNA Technologies).

### Antibodies

*Unlabelled primary antibodies*: anti-human CD8α antibody (Cell Signaling Technology, Cat#85336), anti-human CD28 antibody (Cell Signaling Technology, Cat#38774S), anti-human CD45 antibody (Cell Signaling Technology, Cat#13917S), anti-human PD-1 antibody (Cell Signaling Technology, Cat#86163T), anti-human Lck antibody (Cell Signaling Technology, Cat#2787S), anti-human ZAP-70 antibody (Cell Signaling Technology, Cat#3165S), anti-human LAT antibody (Cell Signaling Technology, Cat#45533S), anti-human PLCγ1 antibody (Cell Signaling Technology, Cat#5690S), and anti-human phospho-ZAP-70 (Tyr319) antibody (Cell Signaling Technology, Cat#2701).

*Labelled secondary antibodies*: goat anti-rabbit secondary antibody Fab AF647 (ThermoFisher, Cat#A66788), goat anti-rabbit secondary antibody F(ab’)2 AF568 (ThermoFisher, Cat#A-21069).

*Labelled primary antibodies*: anti-human TCRαβ BV421 (Biolegend, Cat#306722), anti-human TCR Vβ13.1 PE (Biolegend, Cat#362410), anti-human CD3ε AF488 (OKT3, Biolegend, Cat#317310), anti-human CD28 AF647 (Biolegend, Cat#302953), anti-human CD8 BV711 (Biolegend, Cat#344734), anti-human CD4 AF647 (Biolegend, Cat#300520), anti-human Lck antibody AF647 (Biolegend, Cat#628303), anti-human ZAP-70 antibody AF647 (Biolegend, Cat#693507), and anti-human PD-1 antibody AF488 (Biolegend, Cat#329935).

### Home-made recombinant proteins

#### UCHT-1 Fab

*Sequence:* DIQMTQSPSSLSASVGDRVTITCRASQDIRNYLNWYQQKPGKAPKLLIYYTSRLESGVPSRFSGSGSGTDYTLTISSLQPEDFATYYCQQGNTLPWTFGQGTKVEIKRADAAPTVSIFPPSSEQLTSGGASVVCFLNNFYPKDINVKWKIDGSERQNGVLNSWTDQDSKDSTYSMSSTLTLTKDEYERHNSYTCEATHKTSTSPIVKSFNRNECEVQLVESGGGLVQPGGSLRLSCAASGYSFTGYTMNWVRQAPGKGLEWVALINPYKGVSTYNQKFKDRFTISVDKSKNTAYLQMNSLRAEDTAVYYCARSGYYGDSDWYFDVWGQGTLVTVSSAKTTPPSVYPLAPGSAAQTNSMVTLGCLVKGYFPEPVTVTWNSGSLSSGVHTFPAVLQSDLYTLSSSVTVPSSTWPSETVTCNVAHPASSTKVDKKIVPRDCGCHHHHHHHHHHHH.

#### ICAM-1 extracellular domain

*Sequence:* ETGQTSVSPSKVILPRGGSVLVTCSTSCDQPKLLGIETPLPKKELLLPGNNRKVYELSNVQEDSQPMCYSNCPDGQSTAKTFLTVYWTPERVELAPLPSWQPVGKNLTLRCQVEGGAPRANLTVVLLRGEKELKREPAVGEPAEVTTTVLVRRDHHGANFSCRTELDLRPQGLELFENTSAPYQLQTFVLPATPPQLVSPRVLEVDTQGTVVCSLDRLFPVSEAQVHLALGDQRLNPTVTYGNDSFSAKASVSVTAEDEGTQRLTCAVILGNQSQETLQTVTIYSFPAPNVILTKPEVSEGTEVTVKCEAHPRAKVTLNGVPAQPLGPRAQLLLKATPEDNGRSFSCSATLEVAGQLIHKNQTRELRVLYGPRLDERDCPGNWTWPENSQQTPMCQAWGNPLPELKCLKDGTFPLPIGESVTVTRDLEGTYLCRARSTQGEVTREVTVNVLSPRYECAHHHHHHHHHHHH.

#### CD86 extracellular domain

*Sequence:* ETGAPLKIQAYFNETADLPCQFANSQNQSLSELVVFWQDQENLVLNEVYLGKEKFDSVHSKYMGRTSFDSDSWTLRLHNLQIKDKGLYQCIIHHKKPTGMIRIHQMNSELSVLANFSQPEIVPISNITENVYINLTCSSIHGYPEPKKMSVLLRTKNSTIEYDGIMQKSQDNVTELYDVSISLSVSFPDVTSNMTIFCILETDKTRLLSSPFSIELEDPQPPPDHIPKCAHHHHHHHHHH.

#### PD-L1 extracellular domain

*Sequence:* ETGFTVTVPKDLYVVEYGSNMTIECKFPVEKQLDLAALIVYWEMEDKNIIQFVHGEEDLKVQHSSYRQRARLLKDQLSLGNAALQITDVKLQDAGVYRCMISYGGADYKRTIVKVNAPYNKINQRILVVDPVTSEHELTCQAEGYPKAEVIWTSSDHQVLSGKTTTTNSKREEKLFNVTSTLRINTTTNEIFYCTFRRLDPEENHTAELVIPELPLAHPPNERKCAHHHHHHHHHHHH HHHH.

### Cell lines

Jurkat E6-1 cells were purchased from ATCC (Cat#TIB-152) and cultured in RPMI 1640 culture medium (RPMI 1640 medium supplemented with 10% fetal bovine serum, 1% Penicillin-Streptomycin, and Glutamine). 293T cells were purchased from ATCC (Cat#CRL-3216) and cultured in DMEM culture medium (DMEM medium supplemented with 10% fetal bovine serum, 1% Penicillin-Streptomycin, and Glutamine). Expi293F cells were purchased from ThermoFisher Scientific (Cat#A14527) and cultured in Expi293^TM^ Expression Medium without serum.

### Construction of plasmids

The 1G4 TCR–GFP, CD8αβ, CD28–GFP, Lck–GFP, and ZAP-70–GFP plasmids were generated by separately inserting full-length human 1G4 TCR αβ chains (α-GSG-T2A-β), CD28, Lck, and ZAP-70 cDNAs flanked with C-terminal GFP into the pHR vector with PCR and infusion kits. The CD8 plasmid was generated by inserting full-length human CD8 αβ chains (α-GSG-T2A-β) cDNA into the pHR vector with PCR and infusion kits. All plasmids were verified by sequencing at Source BioScience.

### Recombinant protein expression and purification

The recombinant protein with an N-terminal signaling peptide and a C-terminal 12×His tag was transiently expressed in Expi293F cells. Cells were cultured in suspension and transfected using polyethyleneimine (PEI) according to the manufacturer’s protocol. After incubation for 72 h, cells were discarded, and the supernatant was harvested. The supernatant was then incubated with Ni–NTA affinity resin. The resin was washed with wash buffer (30 mM imidazole in PBS), and the target protein was eluted with elution buffer containing 500 mM imidazole in PBS. The eluted protein was further purified and buffer-exchanged into PBS by gel filtration chromatography.

### Fluorescence labelling of protein

The purified recombined protein solution was mixed with the Alexa Fluor dye (succinimidyl ester) in a molar ratio of 8:1 (dye-to-protein). The total volume was filled to 50 μL with 5 μL sodium bicarbonate (1 M, pH=8.3) and additional PBS buffer. The mixture was kept at 4°C overnight with light blockade. According to the manual instructions, the extra dye was removed from the labelled protein using a desalting column. The F:P ratio was calculated using measurements from the nanodrop2000.

### Gene knockout (KO) with CRISPR Cas9 in Jurkat cells

7.5 µL Cas9 protein (21 μM) was mixed with 2 μL (100 μM) grRNA targeting TCR, CD28, Lck, or ZAP-70, and the mixture was incubated at 37°C for 15 min. After that, 1 µL enhancer (200 µM) was added to finalise the CRISPR–Cas9 mixture. 1.5 million Jurkat cells were harvested and washed with Opti-MEM^TM^ medium twice, and cells were suspended in Opti-MEM^TM^ medium plus CRISPR–Cas9 mixture to 100 µL. Cells were transferred to a 2-mm cuvette for electroporation with the ECM 830 (BTX) using the following parameters: 275 V, 2 ms, and one pulse. After that, cells were cultured in the RPMI 1640 culture medium at 37°C with 5% CO_2_. After 3–5 days, gene expression in cells was verified using flow cytometry with extracellular or intracellular staining. Cell sorting was done if the negative population was lower than 85%. Otherwise, cells were used directly after the KO.

### Preparation of lentivirus and construction of cell lines

2 µg gene-expressing plasmid, 2 µg lentivirus envelope plasmid pMD2.G, and 2 µg lentivirus packaging plasmid p8.91 were mixed into 100 µL Opti-MEM^TM^ medium, and 9 µg PEI was added to 100 µL Opti-MEM^TM^ medium. After 5 min of incubation at RT, PEI solution was added to the gene mixture, and incubated at RT for another 20 min. Gene–PEI mixture was slightly added into 293T cells in a 6-well plate at a 70% cell density. 293T cells were cultured in the DMEM culture medium at 37°C with 5% CO_2_ for 2 days. Then, lentivirus was harvested by collecting the supernatant and concentrated with a 10 kDa cutoff tube. 1.5 million cells were suspended with lentivirus solution and cultured in RPMI 1640 culture medium at 37°C with 5% CO_2_. After 24 h, the virus was removed by washing cells twice, and gene expression in cells was verified using flow cytometry with extracellular or intracellular staining or the fused GFP signal after another day. Cell sorting was done if the positive population was lower than 90%. Otherwise, cells were used directly after the experiment.

### Intensity quantification of the single-molecule tag on the coverslip surface

100 μL of fluorophore-labelled or intrinsically fluorescent proteins in PBS solution with a concentration of around 0.1 ng/mL was injected into the plasma-cleaned glass-bottom channel, incubating at room temperature (RT), 24℃ for 1 h. The protein was treated with 4% PFA for 10 min at RT or remained natural; then, the channel was washed twice with PBS before imaging. A single-molecule image was taken in total internal reflection fluorescence microscopy (TIRFM) with specific laser power (%) and exposure time (ms) at RT or 37°C. The TIRF angles for 405 nm, 488 nm, 561 nm, and 640 nm lasers are 76.56°, 80.61°, 77.95°, and 73.12°, corresponding to a 56 nm-, 62 nm-, 75 nm-, and 100 nm-penetration depth, respectively. The full powers of 488 nm, 561 nm, and 640 nm lasers are 100 mW, 150 mW, 150 mW, and 140 mM, respectively. Single molecules were isolated in 5×5-pixel regions and analysed in Fiji to quantify the mean fluorescent intensity of single molecules and nearby backgrounds. The following formula calculated the fluorescent intensity of a single molecule:

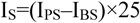

Where I_S_ is the total fluorescent intensity of a single molecule, I_PS_ is the measured mean fluorescent intensity of the protein in the 5×5-pixel region, and I_BS_ is the measured mean fluorescent intensity of the background in the nearby blank 5×5-pixel region. I_PS_–I_BS_ is the mean fluorescent intensity of the single molecule. 25 is the area of the 5×5-pixel region. 16 independent single molecules and backgrounds were analyzed to determine the mean and total fluorescent intensity of this single molecule. All of the single-molecule intensity quantifications were done at RT 24℃ unless stated. The F:P ratio of each protein molecule in this study was maintained above 1.0, ensuring that every single molecule was labelled with at least one fluorophore. The only exception in this study is the dataset presented in Extended Data Fig. 1d and 1g. For molecules with an F:P ratio < 1, the mean intensity was recalibrated by multiplying the measured value by the corresponding F:P ratio. For molecules with an F:P ratio > 1, no recalibration of the mean intensity was performed.

### Intensity quantification of multiple molecule tags on the surface

Using the identical imaging procedures that are used for imaging single molecules, multiple molecules in a region on a surface were imaged and then analyzed in Fiji to quantify the mean fluorescent intensity of the region. The same image of the region without fluorescent molecules, which serves as the background, was imaged and analyzed. The following formula was used to calculate the intensity of multiple molecules in the region:

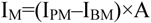

Where I_M_ is the total fluorescent intensity of multiple molecules, I_PM_ is the measured mean fluorescent intensity of proteins in this region, and I_BM_ is the measured mean fluorescent intensity of the background. I_PM_–I_BM_ is the mean fluorescent intensity of these proteins. A is the region’s area, for example, A is 10^4^ for a 100×100-pixel region. Three independent TIRF images of molecules and backgrounds were analyzed to determine the fluorescent intensity of multiple molecules. All of the quantifications were done at RT 24℃ unless stated.

### Quantification of molecular numbers and densities on the SLBs using QuEST

The protocol for preparing the supported lipid bilayers (SLBs) has been published in our previous studies^58,59^. Briefly, the lipid solution was made by mixing DOPC lipid and Ni–NTA lipid in a molar ratio. Unless stated, the nonfluorescent SLBs in this study consist of 12.5% Ni–NTA and 87.5% DOPC for all experiments. 60 μL of lipid solution was injected into the plasma-cleaned glass-bottom channel, incubating for 30 min to form the SLBs on the coverslip. Extra lipid was washed twice with HBSS buffer (20 mM Hepes, 137 mM NaCl, 5 mM KCl, 6 mM D-glucose, 1 mM CaCl_2_, 2 mM MgCl_2_, and 0.1 mg/mL BSA), then the channel was blocked in HBSS buffer at RT for 10 min. After that, 100 μl fluorescence-labelled proteins with His tags in ideal concentrations were incubated in the channel at RT for 1 h. Extra protein was removed by washing the channel three times using HBSS buffer before imaging. After that, fluorescence-labelled proteins on the SLBs were captured in a 550×550-pixel region with TIRFM. After quantifying the intensity of the molecules in this region, the following formula was used to calculate the number of molecules in a region on the SLBs:

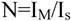

where N is the number of molecules in the region, I_M_ is the total fluorescent intensity of multiple molecules in this region, and I_s_ is the fluorescent intensity of the corresponding single molecule. The method of quantification of I_M_ and I_S_ was introduced above. After that, the following formula was used to calculate the density of molecules in a region on the SLBs:

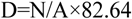

Where D is the density of molecules in the region (unit: molecule/μm^2^), N is the number of molecules in this region, and A is the area of this region. In our imaging setup, one pixel corresponds to 0.11 μm, so 1 μm^2^ is equivalent to 82.64 pixel^2^.

Using the QuEST to calculate regions in different areas follows the same rules, with adjustments only to the region size.

### Quantification of molecular numbers and densities on the SLBs, or on cell surface, or in the TCR signalosome using QuEST

For experiments with cells interacting with the protein-coated SLBs, the molecule numbers and densities in a region were calculated in the same way as described above. The analyzed region on or near the cell surface was determined based on the location of signaling molecules. In Fig. 2 and Extended Data Fig. 8, for Jurkat cells with scanning TCRs, the analyzed regions were chosen based on the location of TCR–GFP, CD28–GFP, Lck–GFP, or ZAP-70–GFP in different scenarios; For Jurkat cells with activated TCRs, the analyzed regions were chosen based on the location of the TCRs. In Fig. 3–5, for both scanning and activated TCRs in T cells, the analyzed regions were chosen based on the location of the TCRs, which were stained by UCHT-1.

### Relative error of molecular quantification of numbers and densities using QuEST

The calculation error arises from the missed fluorescent intensity of molecules near the edge of the measured region. Due to the Abbe diffraction limit in TIRFM, the intensity of a single molecule is distributed over a 5×5-pixel array. When selecting a circular measurement area, approximately 20% of the fluorescence intensity from edge-proximal molecules was lost (grey region, Extended Data Fig. 2a). This fluorescence loss affects all molecules interacting with the boundary. For simplicity, we model the global distribution of these molecules as a single-line distribution, where only those intersecting the dashed grey line (Extended Data Fig. 2a) exhibit intensity loss. Within this group, the fluorescence loss decreases linearly with distance from the edge, from 20% at the boundary to 0% farther away. Consequently, the average fluorescence loss for edge molecules is approximately 10%.

Altogether, the following formula was refined to calculate the relative error in the quantification using the QuEST method:

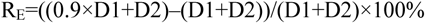

Where R_E_ is the relative error, D1 is the distance of the grey dashed line, and D2 is the distance of the red solid line in Extended Data Fig. 2a.

### Determination of the linear increase range of fluorescent protein density

The SLBs consisting of nonfluorescent DOPC and fluorescent DOPE in varied concentrations were imaged in TIRFM to determine the linear increase ranges of different fluorescent DOPE lipids. The linear increase range of density of different fluorescent DOPE lipids was therefore calculated with the parameter of the diameter of the DOPE lipid head (0.8 nm). The linear increase range of fluorescent molecule density was transferred from the DOPE lipid range based on their molecular diameters (UCHT-1 Fab: 3 nm, ICAM-1: 2 nm, GFP: 3 nm, PD-L1: 2.2 nm, and CD86: 2.4 nm). The molecular diameters were measured based on the molecular structures displayed in Chimera. The quantified molecular densities of the molecules in this study were located in their linear increase ranges, which excludes the possibility of self-quenching between fluorophores.

### Isolation and culturing of T cells

Human peripheral blood samples were obtained under Human Tissue Authority (HTA) licence 12217 at the John Radcliffe Hospital, University of Oxford. The study was approved by the NHS Health Research Authority (REC reference 11/H0711/7; IRAS ID 66954), and all donors provided written informed consent. All sample procedures complied with the national guidelines and regulations. CD8^+^ T cells were isolated from human blood using the human CD8^+^ T cell enrichment cocktail according to manual instructions. A million/mL primary CD8^+^ T cells were pre-activated by incubating with the same density of anti-CD3/CD28 magnetic beads in the RPMI 1640 culture medium with 50 units/mL human IL-2 for 2 days. From this point onward, primary CD8⁺ T cells were converted into CD8⁺ T cell blasts, which exhibit greater protein homogeneity across different T cell subsets. After that, beads were removed, and CD8⁺ T cell blasts were maintained in the RPMI 1640 culture medium with 50 units/mL human IL-2 for another 3 days before use. CD8⁺ T cell blasts were kept for 10 days for experiments after bead removal. Unless otherwise stated, the T cells used in this study were CD8^+^ T cell blasts generated with this method.

### Interaction between cells and protein-functionalized SLBs

For the scenario of scanning TCRs, 100 μl protein solution of 0.1 μg/mL ICAM-1 was incubated on the SLBs in the channel at RT for 1 h to have the molecular densities of 177 ICAM-1/μm^2^. For the scenario of activated TCRs, 100 μl protein solution of 0.02 μg/mL UCHT-1 and 0.1 μg/mL ICAM-1 was used to incubate to have the molecular densities of 36 UCHT-1/μm^2^ and 177 ICAM-1/μm^2^. If inhibitory or costimulatory signaling is required, additional CD86 (0.387 μg/mL) or PD-L1 (1 μg/mL) in a density of 400 molecules/μm^2^ were integrated onto the SLBs with UCHT-1 and ICAM-1. Half a million T cells or Jurkat cells were washed twice with HBSS buffer and then injected into the channel to incubate with the protein-functionalized SLBs at 37°C. 20-minute and 25-minute incubating times were used for T cells and Jurkat cells, respectively, in this study unless stated. Next, cells were treated with the BD fixation/permeabilization kit and incubated with primary antibody (diluted 200 times) at 4°C overnight. The next day, cells were incubated with goat anti-rabbit secondary Fab AF647 (diluted 200 times) at RT for 1.5 h before TIRF imaging. Cells not requiring intracellular staining were imaged after fixation of cells. Cells were imaged at RT unless stated. Cells were imaged at 37°C directly without fixation and permeabilization for live cell imaging.

### Quantification of the molecular loss after cell treatments

The Jurkat cells expressing 1G4 TCR–GFP, CD28–GFP, Lck–GFP, or ZAP-70–GFP were interacted with the protein-functionalized SLBs according to the methods above. Before cell fixation, permeabilization, and staining (FPS), live cells were imaged in the TIRFM, and GFP intensities were acquired. The copy numbers of GFP in live cells were quantified according to the intensity of the single natural GFP proteins using the above methods. After the FPS, the same quantifications on GFP copy numbers were performed according to the intensity of the single PFA-treated GFP proteins. The molecular loss after cell treatments was therefore calculated by comparing values obtained in live and FPS cells. The exact numbers of GFP in both live and fixed cells were calculated considering the 30% dark proportion factorof GFP in cells^33^.

### Quantification of the UCHT-1 to TCR ratio

#### For scanning TCRs

Half a million 1G4–GFP Jurkat cells were washed twice with HBSS buffer and then injected into the ICAM-1-coated SLBs for a 25-min incubation at 37°C. Cells were then treated with the BD fixation/permeabilization kit. Cells were then washed twice with the BD fixation/permeabilization buffer with the additional 100 mM imidazole to block the interaction of UCHT-1 with the SLBs. From this time point, fixation/permeabilization buffer with the additional 100 mM imidazole was used for the subsequent staining and imaging. Next, TCRs on the cells were incubated with 10 μg/mL UCHT-1 AF568 at RT for 1 h, and extra UCHT-1 was removed before the TIRF imaging. Copy number of TCR was quantified through GFP signal according to the intensity of the single PFA-treated GFP proteins, and the copy number of UCHT-1 was quantified according to the intensity of the single UCHT-1 AF568 proteins. The UCHT-1 to TCR ratio was calculated based on the copy numbers of UCHT-1 and TCR.

#### For activated TCRs

Half a million 1G4–GFP Jurkat cells were washed twice with HBSS buffer and then injected into the UCHT-1 AF568 & ICAM-1-coated SLBs (activated TCRs) for a 25-min incubation at 37°C. Cells were then treated with the BD fixation/permeabilization kit. Copy number of TCR was quantified through GFP signal according to the intensity of the single PFA-treated GFP proteins, and the copy number of UCHT-1 was quantified according to the intensity of the single UCHT-1 AF568 proteins. The UCHT-1 to TCR ratio was calculated based on the copy numbers of UCHT-1 and TCR.

### Quantification of the secondary Fab to signaling molecule ratio

Half a million CD28–GFP, Lck–GFP, or ZAP-70–GFP Jurkat cells were washed twice with HBSS buffer and then injected into the ICAM-1-coated SLBs (scanning TCRs) or UCHT-1 AF568 & ICAM-1-coated SLBs (activated TCRs) for a 25-min incubation at 37°C. Next, cells were treated with the BD fixation/permeabilization kit and incubated with primary antibody (diluted 200 times) at 4°C overnight. The next day, cells were incubated with goat anti-rabbit secondary Fab AF647 (diluted 200 times) at RT for 1.5 h before TIRF imaging. Copy number of CD28, Lck, or ZAP-70 was quantified through GFP signal according to the intensity of the single PFA-treated GFP proteins, and the copy number of secondary Fab was quantified according to the intensity of the single Fab AF647 molecules. The Fab to molecule ratio was calculated based on the copy numbers of secondary Fab and the signaling molecule.

### Mapping the stoichiometry of the TCR signalosome

After the interaction of T cells with the protein-functionalized SLBs, cells were then treated and stained following the above methods.

The density of TCR after cell treatments was acquired by transferring from the number of UCHT-1 using the following formula:

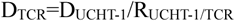

Where D_TCR_ is the density of TCR in cells after treatments, D_UCHT-1_ is the density of UCHT-1, and R_UCHT-1/TCR_ is the UCHT-1 to TCR ratio. For scanning T cells, R_UCHT-1/TCR_ is 0.344; For activated T cells, R_UCHT-1/TCR_ is 0.750 (Fig. 2j).

After that, the following formula was refined to calculate the real density of TCR in cells before any cell treatments:

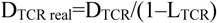

Where D_TCR real_ is the real density of TCR in cells before treatments, L_TCR_ is the TCR loss during cell treatments. For scanning T cells, L_TCR_ is ∼30%; For activated T cells, L_TCR_ is ∼20% (Fig. 2i).

Following the above calculation processes, the density of signaling molecules after cell treatments was acquired by transferring from the number of secondary Fab using the following formula:

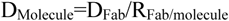

Where D_Molecule_ is the density of signaling molecule in cells after treatments, D_Fab_ is the density of secondary Fab, and R_Fab/molecule_ is the Fab to signaling molecule ratio. For both scanning and activated T cells, R_Fab/molecule_ is ∼1 (Fig. 2j).

After that, the following formula was refined to calculate the real density of signaling molecules in cells before any cell treatments:

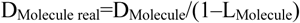

Where D_Molecule real_ is the real density of signaling molecules in cells before treatments, L_Molecule_ is the signaling molecule loss during cell treatments. For both scanning and activated T cells, L_Molecule_ is ∼60% (Fig. 2i).

Stoichiometry of the TCR signalosome was mapped by combining the molecular ratios of eight signaling molecules, including CD8, CD28, CD45, PD-1, Lck, ZAP-70, LAT, and PLCγ1, and the phosphorylated ZAP-70, to the TCR.

### dSTORM imaging

The steps of preparing cell samples on the SLBs and cell treatment were the same as described above, except that the secondary antibodies were changed to goat anti-rabbit F(ab’)2 AF568. Before the dSTORM imaging, the buffer in the sample was changed to the following: 30 mM Tris-HCl, 10 mM NaCl, 5% D-glucose (v/w), 50 mM cysteamine, 0.5 mg/mL glucose oxidase, pH 7.4, to increase the blinking, but to decrease the bleaching of the fluorescent dyes. At the beginning, a 488-nm laser of 100% power was on to excite the UCHT-1 AF488 signal and switch it to a dark state in a 10-ms exposure time; 3000 TIRF images of the UCHT-1 signal were collected in 1 min to bleach most single molecules while avoiding significant focus drifting. Then, a 561-nm laser of 100% power was on to excite the secondary F(ab’)2 antibody signal and switch it to a dark state in a 10-ms exposure time; 6000–9000 TIRF images of the secondary antibody signal were collected in 2–3 min to bleach most single molecules while avoiding significant focus drifting. The images were reconstructed with the ThunderSTORM plugin in Fiji software^60^, and dSTORM images were presented as a form of normalised Gaussian.

### Minimization of FRET effect in the QuEST method

To minimize FRET in the QuEST- method, we used AF488 (or GFP) and AF647 as the fluorescent pair in most imaging experiments, thereby avoiding spectral overlap between AF488 (or GFP) emission and AF647 excitation. In Fig. 2**a**,**c**,**g** and Extended Data Fig. 8**a**,**b**,**d**, we instead used GFP and AF568, or GFP, AF568, and AF647. In these cases, FRET was minimized by the large physical separation between GFP and AF568 or between AF568 and AF647, and was further suppressed by the physical barrier imposed by the cell membrane. In Fig. 3**m**, although AF488 and AF568 were used for dSTORM imaging, FRET was likewise suppressed by the cell membrane, and these measurements were not used for quantification.

## Extended Data Figures and legends

**Extended Data Fig. 1| (Related to Fig. 1).**
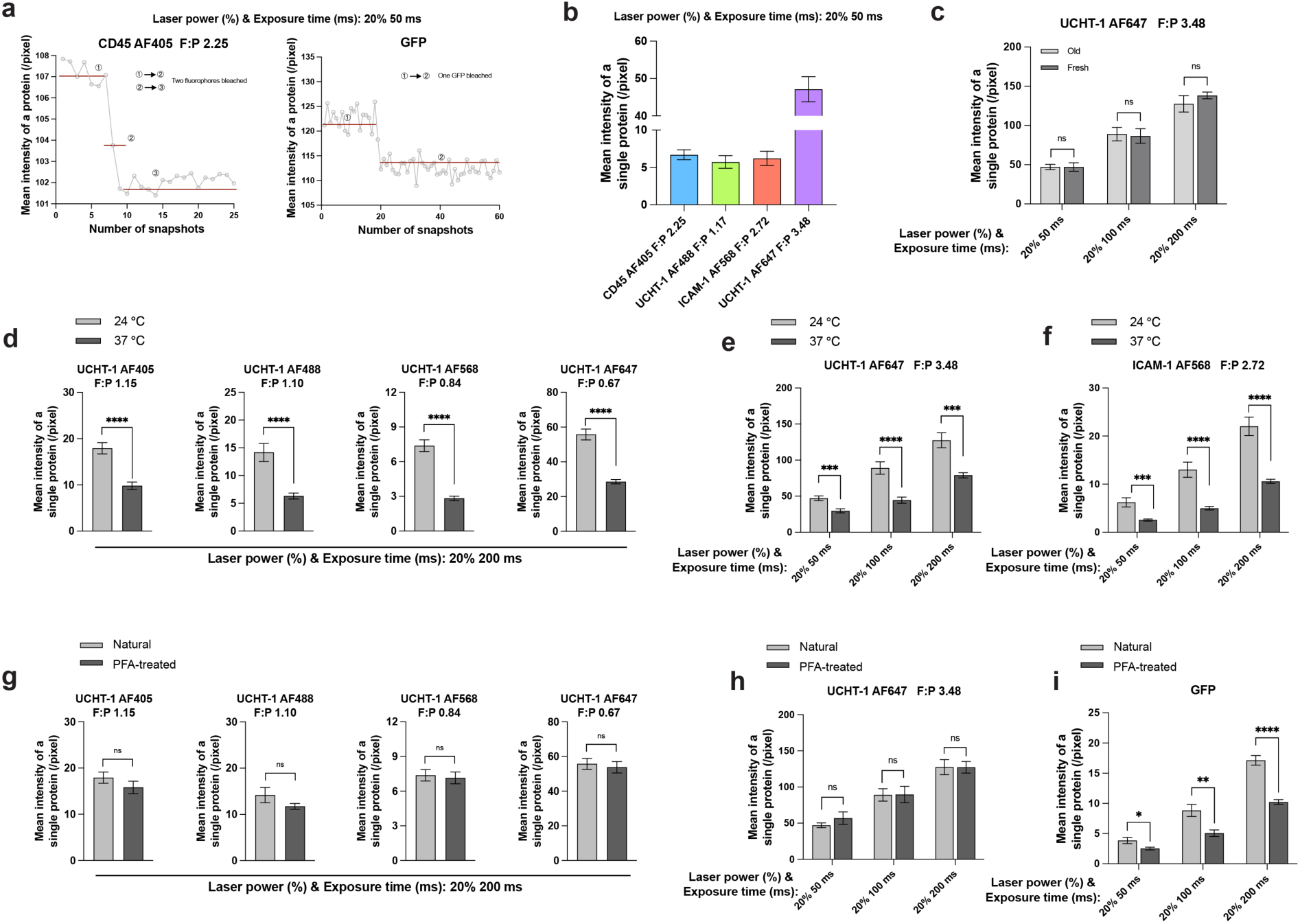
Quantification of fluorescence intensities of single protein molecules under different conditions. **a**, Stepwise photobleaching during repeated TIRF imaging cycles confirms that each fluorescent spot corresponds to a single molecule. **b**, Comparison of mean fluorescence intensities of different single fluorescently labelled protein molecules measured under identical imaging conditions. **c**, Mean fluorescence intensity of single UCHT-1 AF647 molecules at different freshness states. **d**, Mean fluorescence intensities of single UCHT-1 AF405, UCHT-1 AF488, UCHT-1 AF568 and UCHT-1 AF647 molecules measured at different temperatures under identical imaging conditions. **e**, Mean fluorescence intensity of single UCHT-1 AF647 molecules measured at different temperatures under distinct imaging conditions. **f**, Mean fluorescence intensity of single ICAM-1 AF568 molecules measured at different temperatures under distinct imaging conditions. **g**, Mean fluorescence intensities of single UCHT-1 AF405, UCHT-1 AF488, UCHT-1 AF568 and UCHT-1 AF647 molecules following different denaturation treatments under identical imaging conditions. **h**, Mean fluorescence intensity of single UCHT-1 AF647 molecules following different denaturation treatments under distinct imaging conditions. **i**, Mean fluorescence intensity of single GFP molecules following different denaturation treatments under distinct imaging conditions. Unless otherwise indicated, measurements were performed at 24 °C. Bar plots indicate the mean (bar height) and 95% confidence interval (error bars), with individual data points overlaid for each bar. Data are mean ± SEM. Statistical significance was assessed using an unpaired two-tailed Student’s *t*-test (\*\*\*\**P* ≤ 0.0001; \*\*\**P* ≤ 0.001; \*\**P* ≤ 0.01; \**P* ≤ 0.05; ns, not significant). For **a**, the data are representative of 10 individual single molecules. For **b**–**i**, the data were obtained from 16 individual single molecules.

**Extended Data Fig. 2| (Related to Fig. 1).**
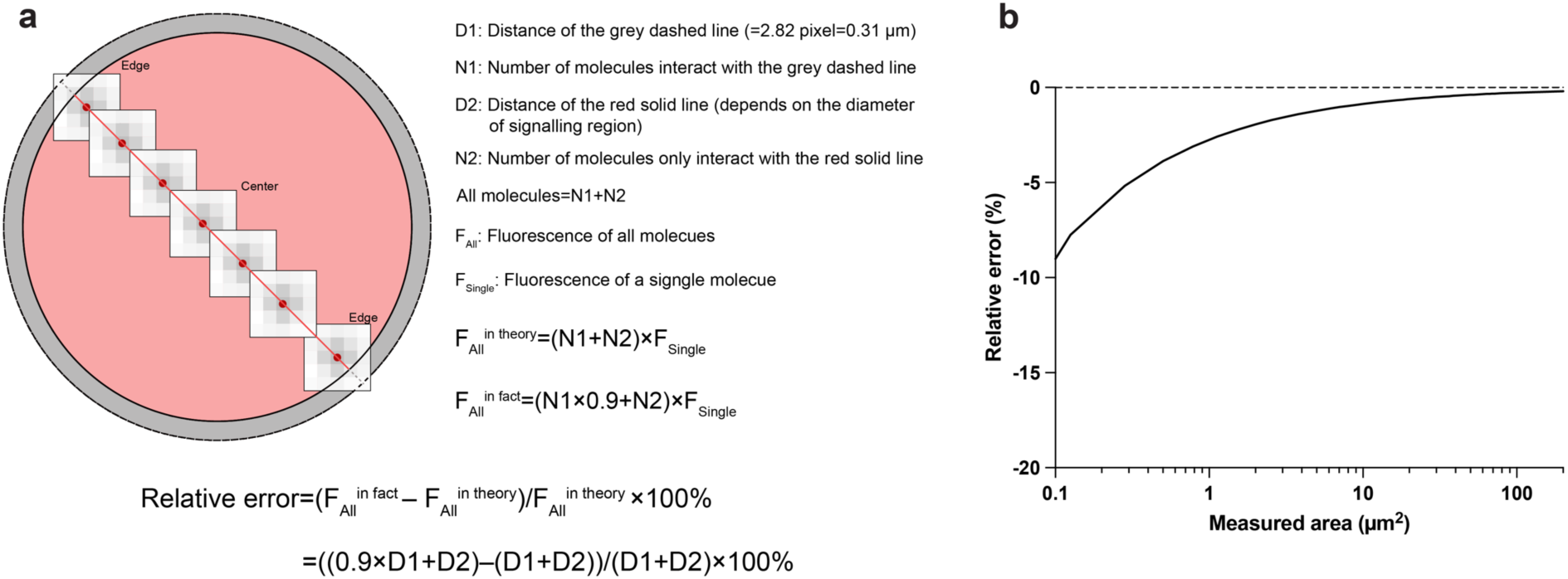
Relative error of molecule density quantification using QuEST. **a**,**b**, Schematic illustration (**a**) and corresponding analysis results (**b**) of relative error estimation for molecule density quantification. The relative error of calculation depends on the diameter of the measured signaling region.

**Extended Data Fig. 3| (Related to Fig. 1).**
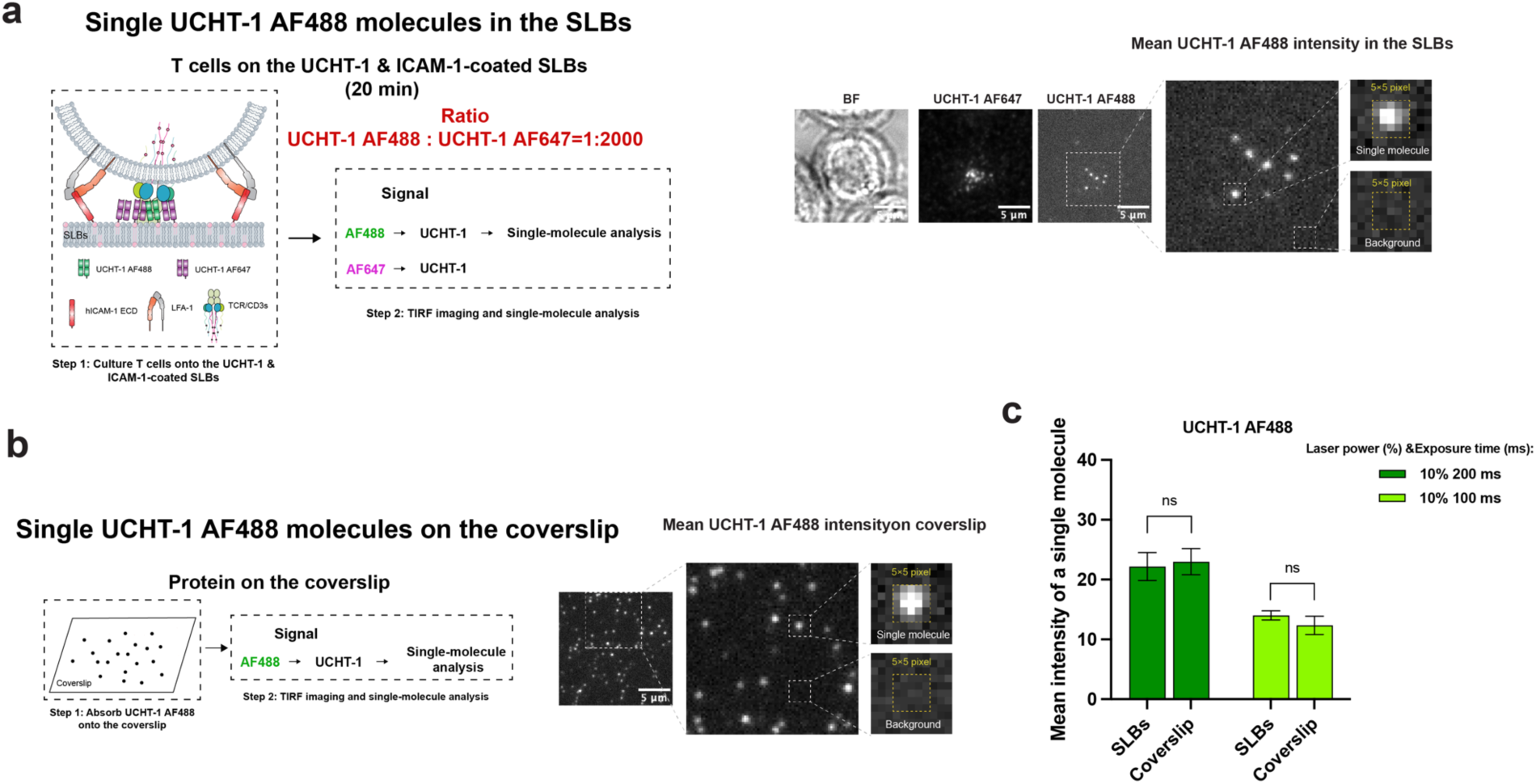
No significant variance in mean fluorescence intensity of single protein molecules quantified on supported lipid bilayers (SLBs) and coverslip. **a**, Workflow (left) and representative TIRF images (right) for quantification of the mean fluorescence intensity of single UCHT-1 molecules on SLBs. **b**, Workflow (left) and representative TIRF images (right) for quantification of the mean fluorescence intensity of single UCHT-1 molecules on coverslip. **c**, Comparison of mean fluorescence intensities measured on SLBs and coverslip. All measurements were performed at 24 °C. Data are presented as mean ± SEM. Statistical significance was assessed using an unpaired two-tailed Student’s *t*-test (ns, not significant). For **c**, data were obtained from 16 single molecules.

**Extended Data Fig. 4| (Related to Fig. 1).**
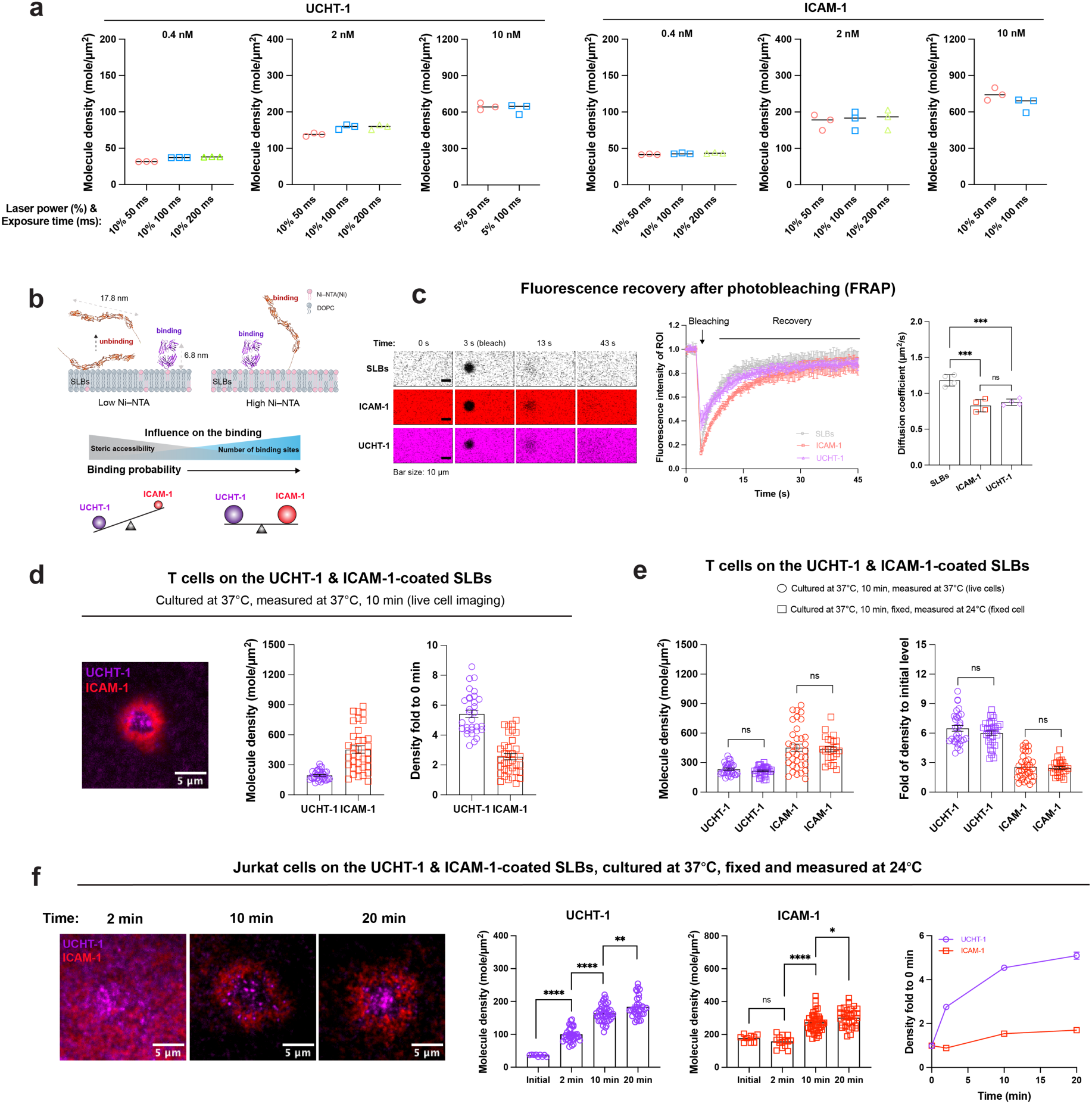
Quantification of molecule numbers and densities in SLBs using QuEST. **a**, Quantified densities of UCHT-1 and ICAM-1 on SLBs under different imaging conditions using the QuEST method. **b**, Schematic illustrating steric accessibility- and binding site-mediated binding of UCHT-1 and ICAM-1 to SLBs. **c**, Fluorescence recovery after photobleaching (FRAP) of UCHT-1 and ICAM-1 on SLBs and quantification of molecular and bilayer fluidity. **d**, Live cell imaging of T cells clustering UCHT-1 and ICAM-1 on SLBs; molecule densities were quantified at 37 °C at the 10-min time point. **e**, Quantified densities of UCHT-1 and ICAM-1 on SLBs measured at different temperatures. **f**, Jurkat cells clustering UCHT-1 and ICAM-1 on SLBs; molecule densities were quantified at different incubation times. Data in **c** are presented as mean ± SD; data in **d**–**f** are presented as mean ± SEM. Statistical significance was assessed using an unpaired two-tailed Student’s *t*-test (\*\*\*\**P* ≤ 0.0001; \*\*\**P* ≤ 0.001; \*\**P* ≤ 0.01; \**P* ≤ 0.05; ns, not significant). Data in **d** and **e** are representative of three independent replicates using T cells from two blood donors. The structural model of UCHT-1 was derived from PDB 3FZU. The structural model of ICAM-1 was generated using AlphaFold3 from the corresponding protein sequences.

**Extended Data Fig. 5| (Related to Fig. 1).**
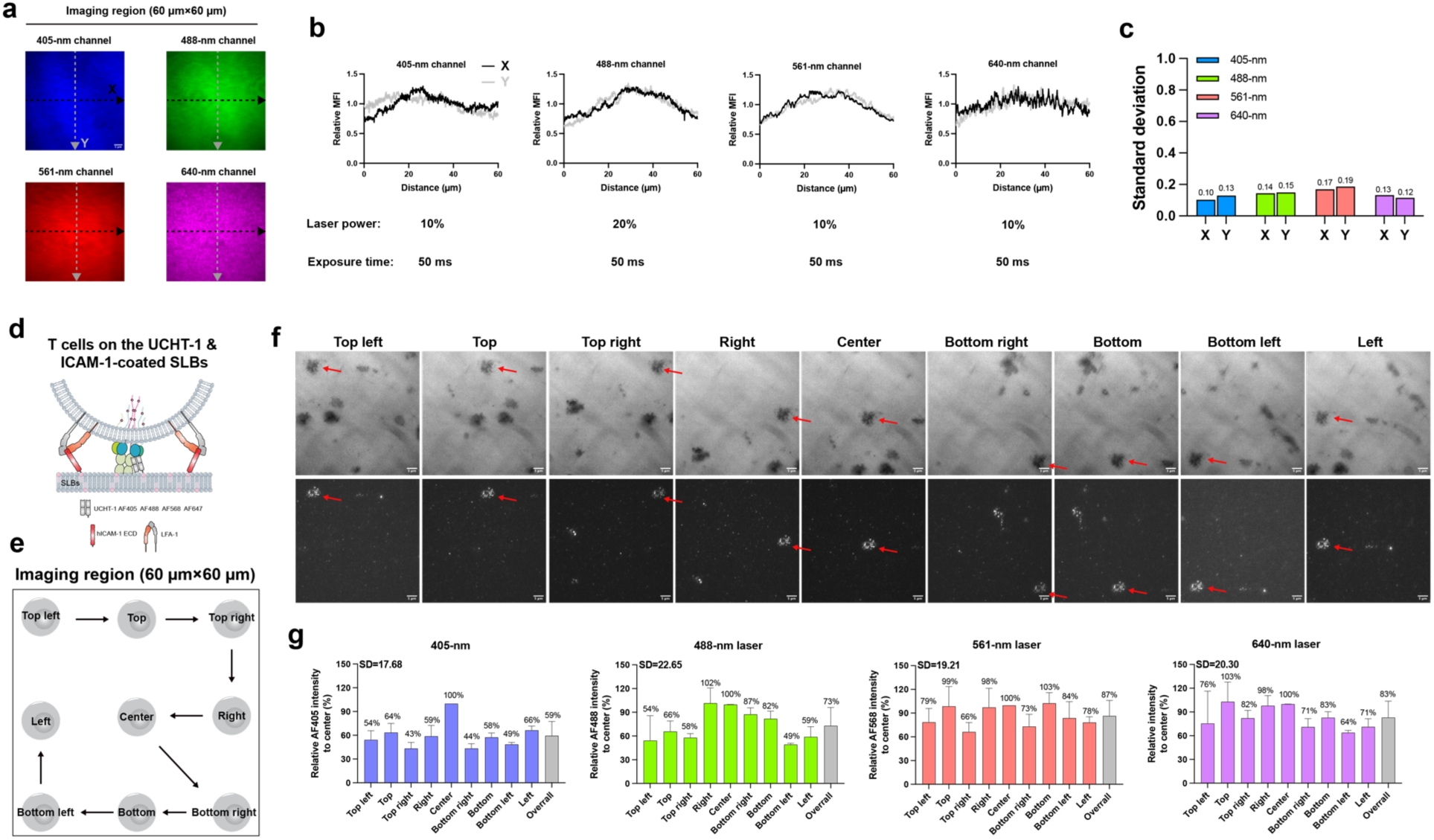
Two-dimensional (2D) uniformity of total internal reflection fluorescence microscopy (TIRFM). **a**, Images of the supported lipid bilayers (SLBs) composed of 1% of each fluorescent DOPE (Atto 390, Atto 488, Atto 565, and Atto 647) and 96% DOPC, captured using fluorescence channels at 405 nm, 488 nm, 561 nm, and 640 nm. X and Y plot-profile positions are marked by dashed arrows. The final image was generated by averaging four independent images. **b**, Mean fluorescence intensities (MFI) along the plot profiles obtained under laser powers with 50-ms exposure times; fluorescence intensities are relative to the average. **c**, Standard deviations (SD) of relative MFI in all channels. **d**, Schematic of the experimental setup. **e**,**f**, Imaging sequence (**e**) and representative images (**f**) of the same sample acquired at different positions within the field of view. **g**, Relative fluorescence intensity at each position normalized to the center of the field of view across four imaging channels. Data are from four independent replicates and presented as mean ± SD.

**Extended Data Fig. 6| (Related to Fig. 2).**
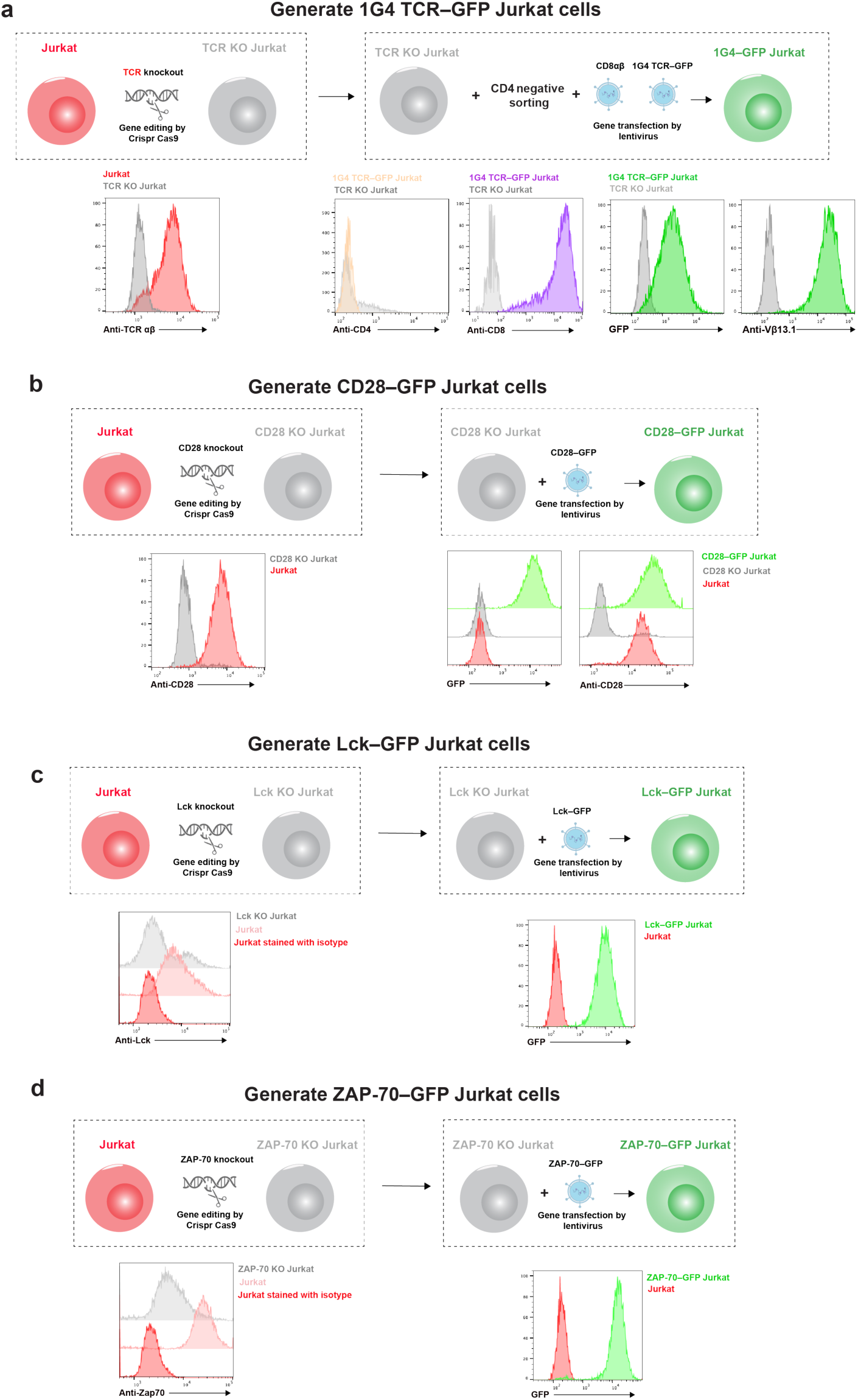
Generation and validation of molecule–GFP Jurkat cells. **a**, Generation (top) and validation (bottom) of 1G4 TCR–GFP Jurkat cells. 1G4 TCR–GFP and CD8 were coexpressed, CD4 was eliminated by negative sorting. **b**, Generation (top) and validation (bottom) of CD28–GFP Jurkat cells. **c**, Generation (top) and validation (bottom) of Lck–GFP Jurkat cells. **d**, Generation (top) and validation (bottom) of ZAP-70–GFP Jurkat cells.

**Extended Data Fig. 7| (Related to Fig. 2).**
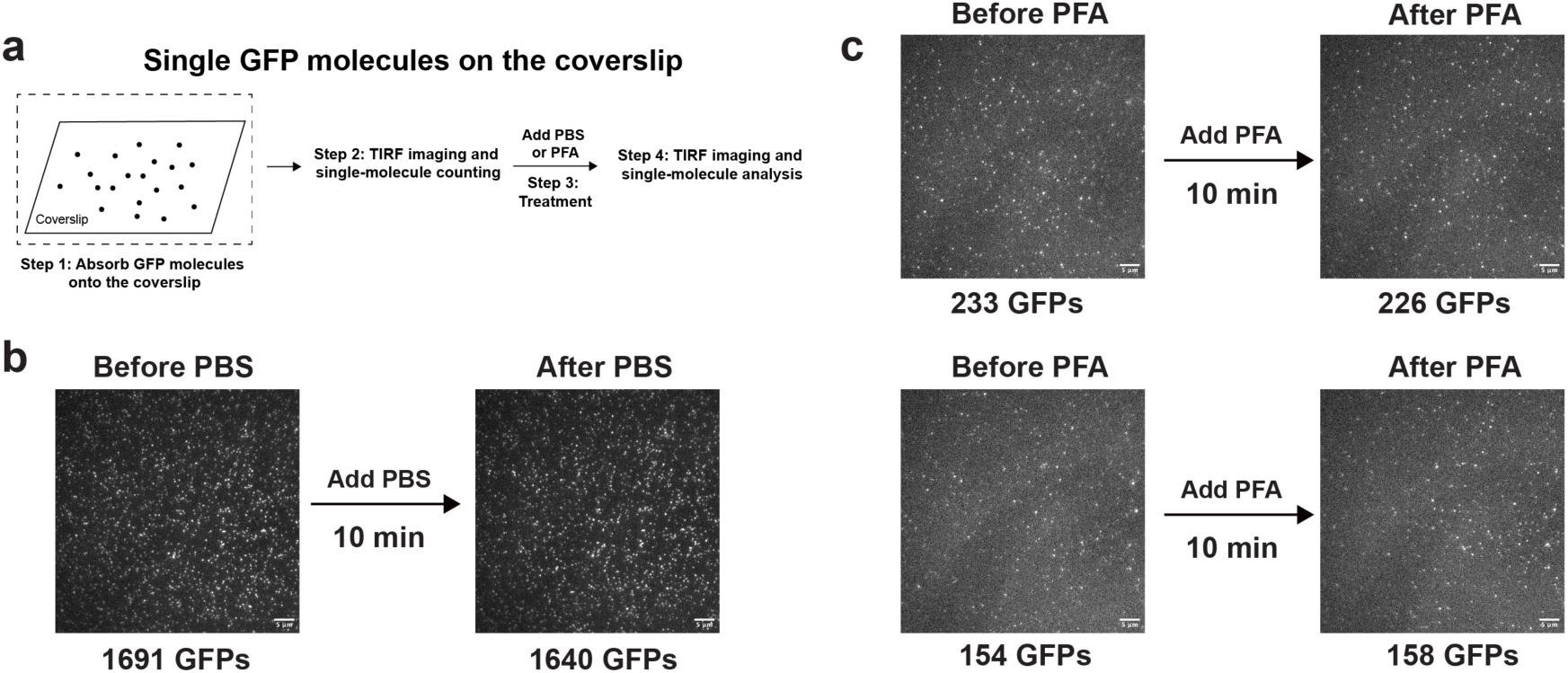
Quantification of GFP molecule numbers before and after PBS or PFA treatment. **a**, Schematic of the experimental design. **b**, Number of GFP molecules quantified before and after PBS treatment. **c**, Number of GFP molecules quantified before and after PFA treatment. The number of single GFP molecules was counted using the TrackMate plugin in Fiji.

**Extended Data Fig. 8| (Related to Fig. 2).**
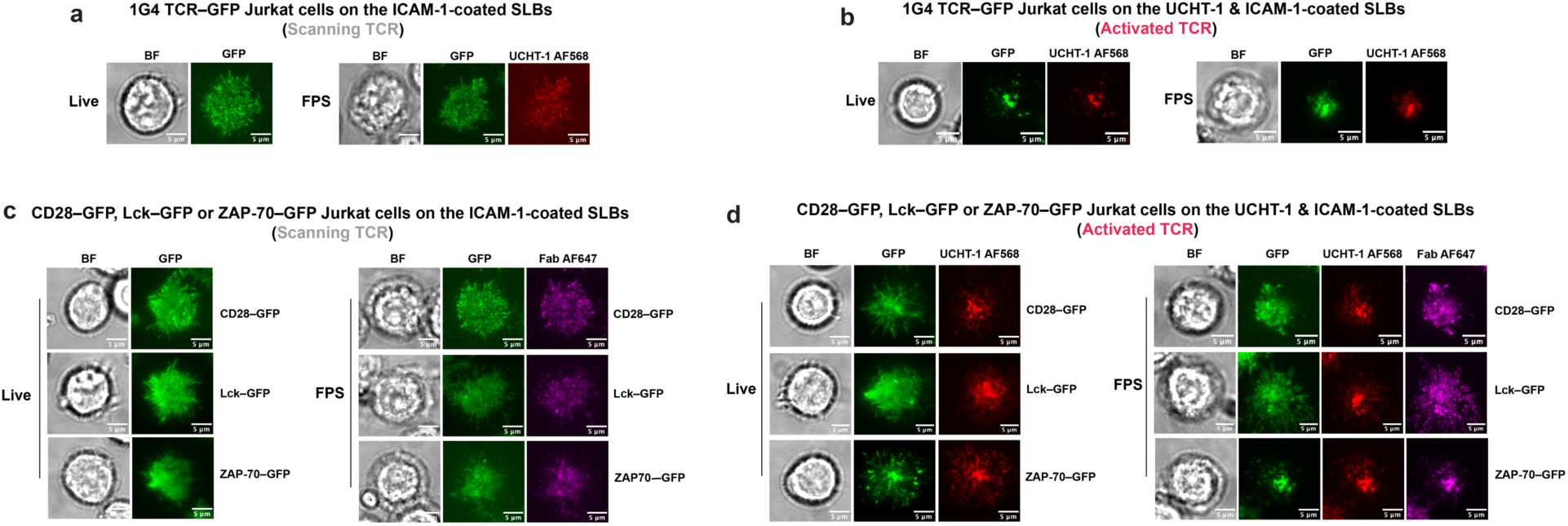
Representative TIRF images for quantification of TCR, CD28, Lck or ZAP-70 loss during cellular treatments and determination of UCHT-1/TCR or Fab/molecule ratio. **a**, Representative TIRF images of 1G4 TCR–GFP Jurkat cells during T cell scanning before and after cell treatments. **b**, Representative TIRF images of TCR–GFP Jurkat cells upon TCR activation before and after cell treatments. **c**, Representative TIRF images of CD28–GFP, Lck–GFP, or ZAP-70–GFP Jurkat cells during T cell scanning before and after cell treatments. **d**, Representative TIRF images of CD28–GFP, Lck–GFP, or ZAP-70–GFP Jurkat cells upon TCR activation before and after cell treatments.

**Extended Data Fig. 9| (Related to Fig. 2).**
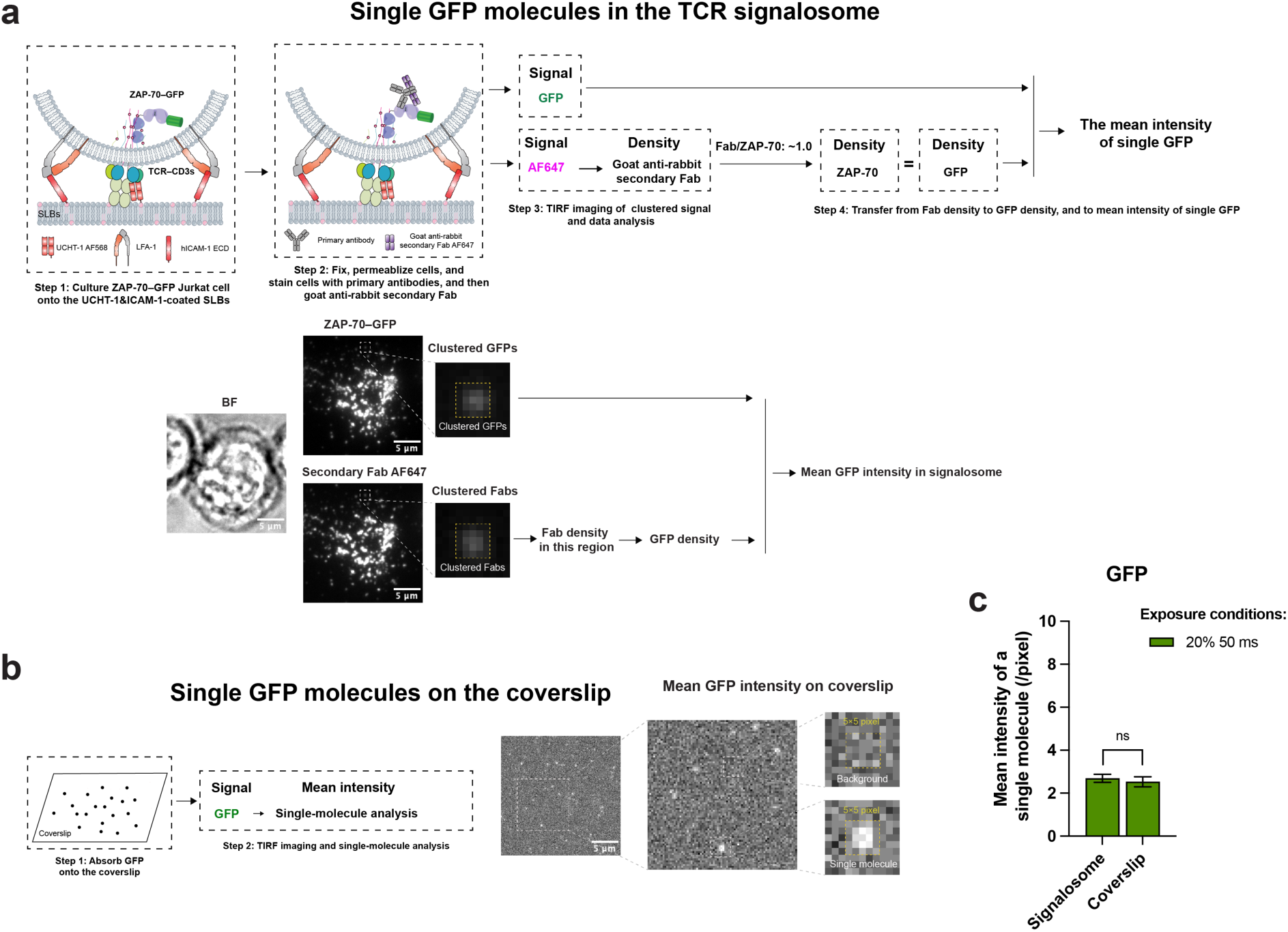
No significant variance in mean fluorescence intensity of GFP quantified in the TCR signalosome and on coverslip. **a**, Workflow (top) and representative TIRF images (bottom) for quantification of the mean fluorescence intensity of single GFP molecules within the TCR signalosome of ZAP-70–GFP expressing Jurkat cells. **b**, Workflow (left) and representative TIRF images (right) for quantification of the mean fluorescence intensity of single GFP molecules immobilized on the coverslip; GFP was fixed with PFA before measurement. **c**, Comparison of mean fluorescence intensities measured in the signalosome and on the coverslip. All measurements were performed at 24 °C. Data are presented as mean ± SEM. Statistical significance was assessed using an unpaired two-tailed Student’s *t*-test (ns, not significant). For **c**, data were obtained from at least 16 individual GFP molecules.

**Extended Data Fig. 10| (Related to Fig. 2).**
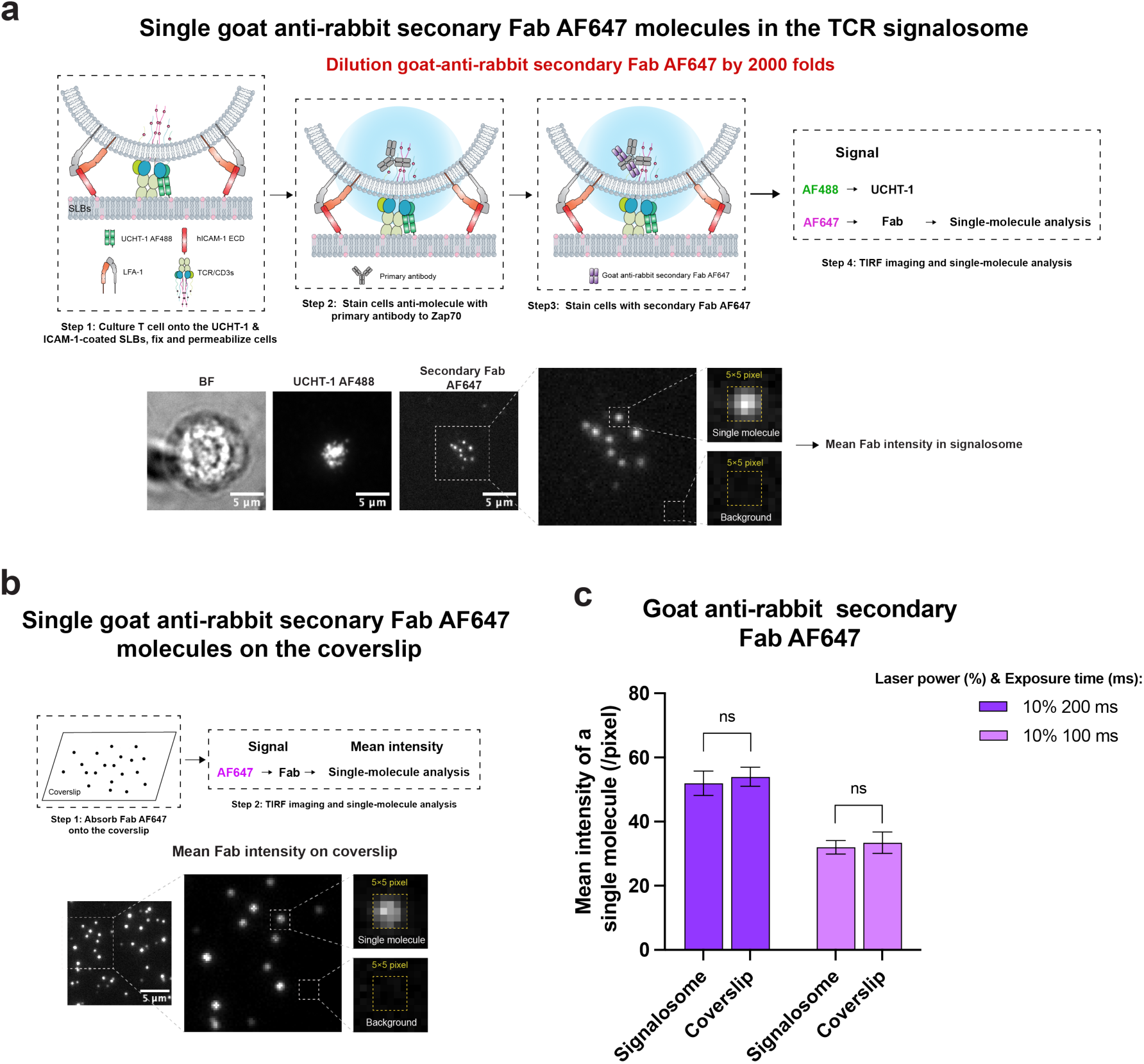
No significant difference in mean fluorescence intensity of single secondary Fab molecules quantified in the TCR signalosome and on coverslip. **a**, Workflow (top) and representative TIRF images (bottom) for quantification of the mean fluorescence intensity of single Fab molecules within the TCR signalosome. **b**, Workflow (top) and representative TIRF images (bottom) for quantification of the mean fluorescence intensity of single Fab molecules immobilized on coverslips. **c**, Comparison of mean fluorescence intensities measured in the signalosome and on the coverslip. All measurements were performed at 24 °C. Data are presented as mean ± SEM. Statistical significance was assessed using an unpaired two-tailed Student’s *t*-test (ns, not significant). For **c**, data were obtained from 16 single molecules.

**Extended Data Fig. 11| (Related to Fig. 2).**
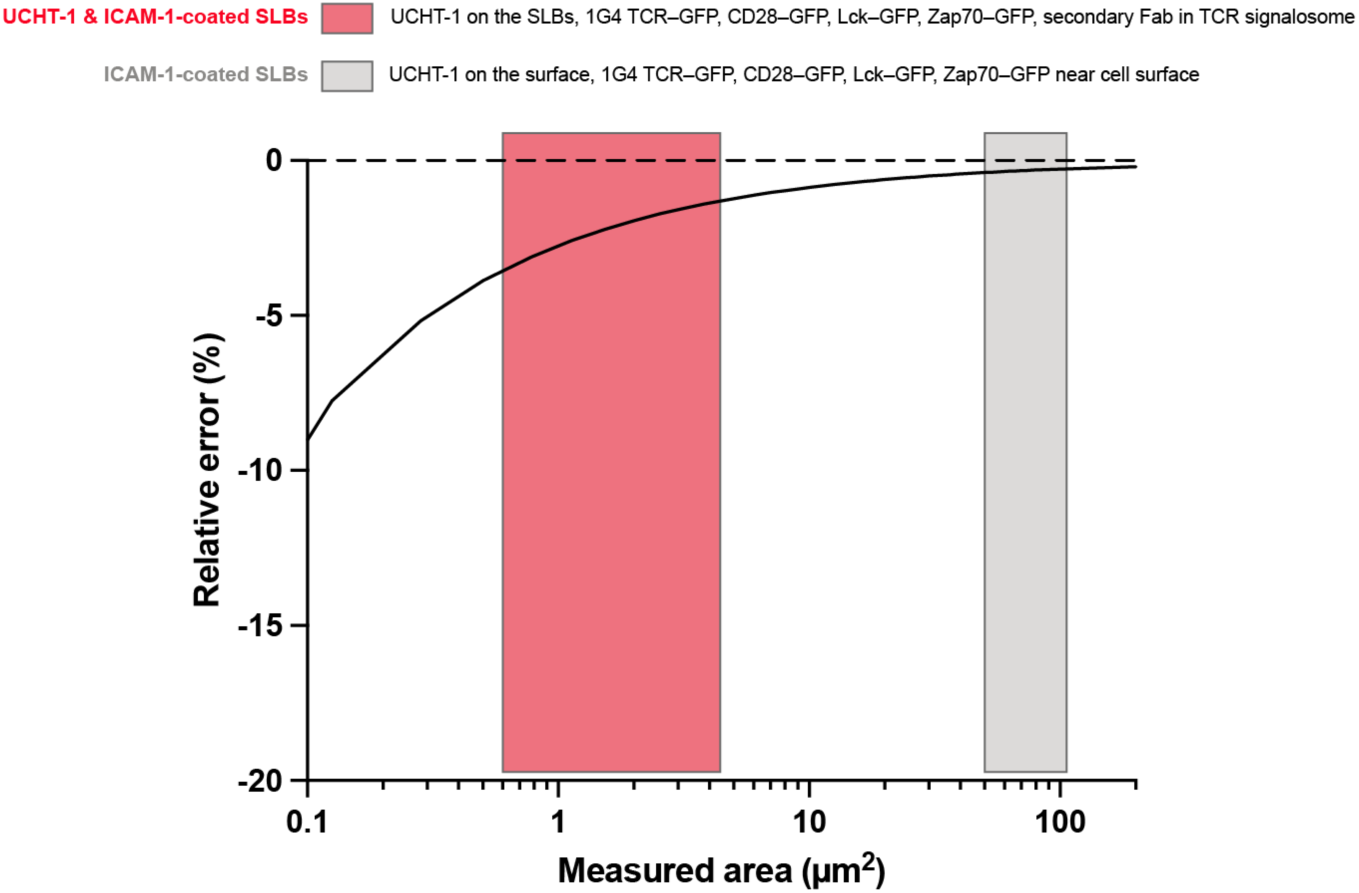
Relative error of molecular quantification of the numbers and densities on the surface or in Jurkat cells at different activation states using QuEST.

**Extended Data Fig. 12| (Related to Fig. 3).**
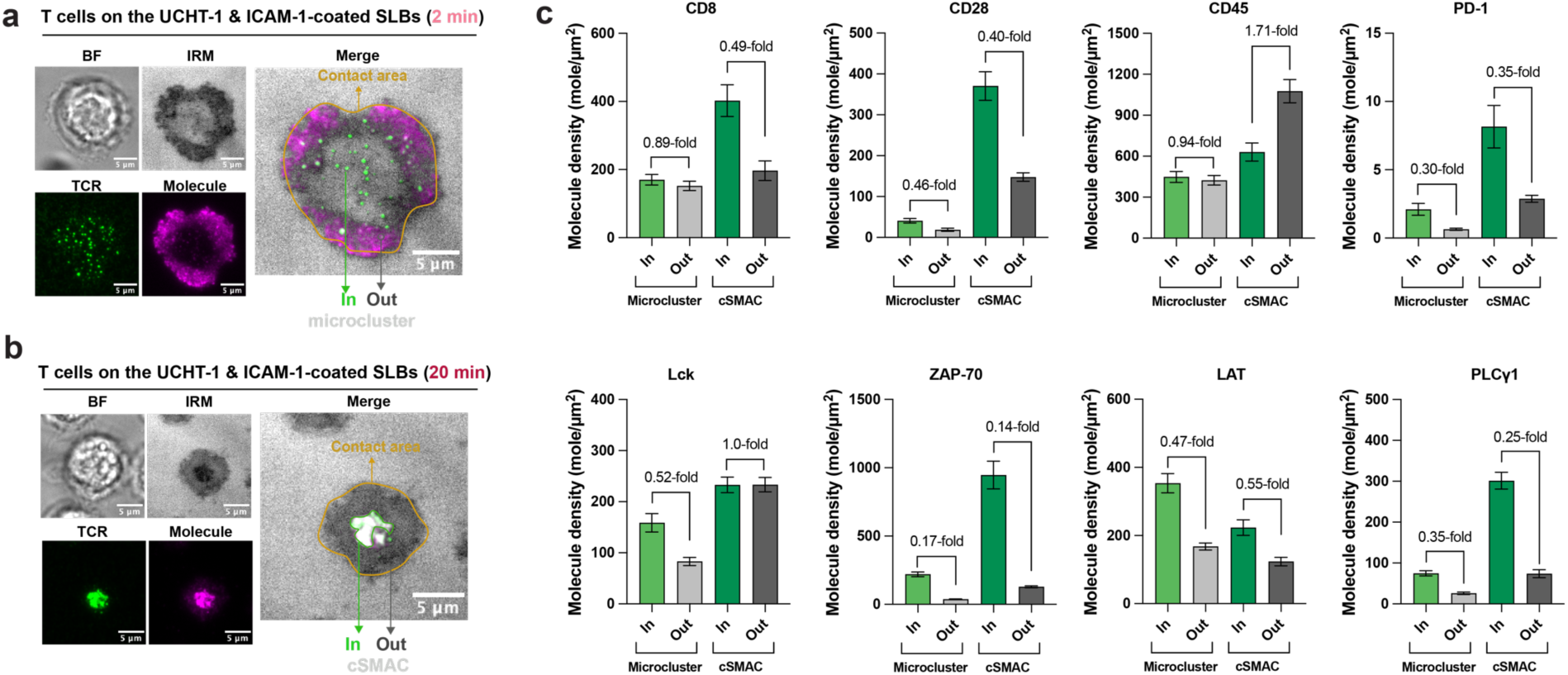
Densities of molecules in and out of the TCR-enriched regions. **a**, Schematic images illustrating the regions analyzed inside and outside microclusters at the early activation stage. **b**, Schematic images illustrating the regions analyzed inside and outside cSMAC at the sustained activation stage. c, Densities of molecules in and out of the TCR-enriched regions. Data are presented as mean ± SEM. At least 30 individual cells were analyzed per condition. Data are representative of three replicates using T cells from three blood donors.

**Extended Data Fig. 13| (Related to Fig. 3).**
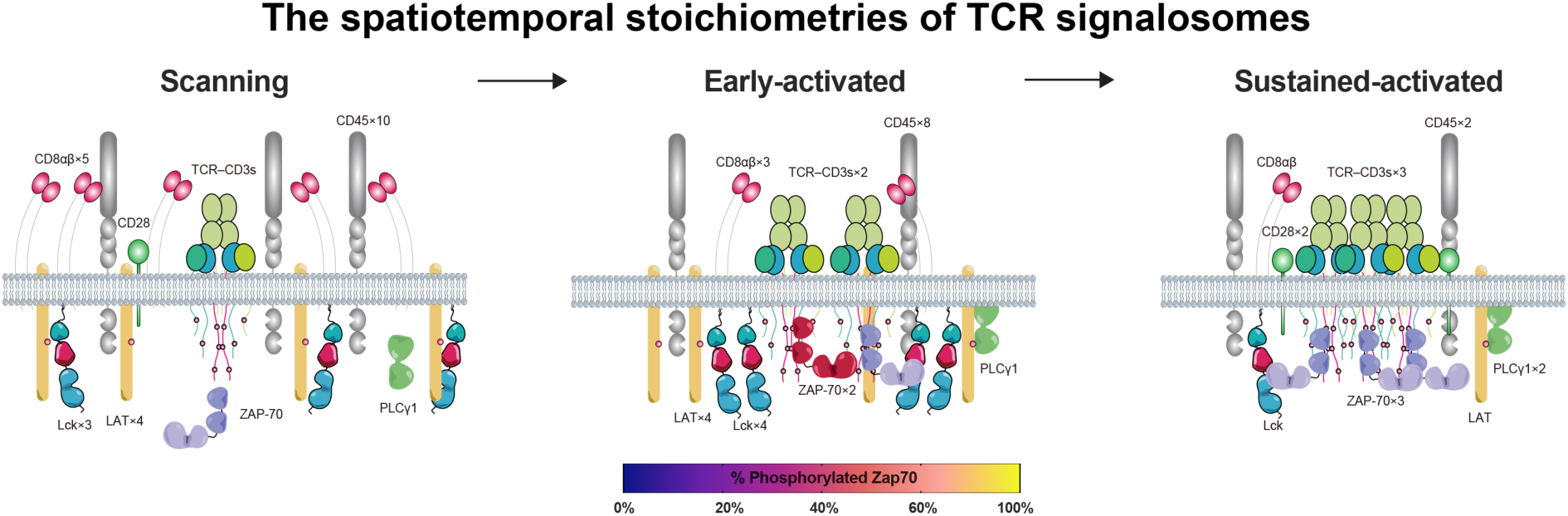
Schematic of the dynamic spatiotemporal stoichiometries of the TCR signalosome with time-dependent ZAP-70 phosphorylation indicated.

**Extended Data Fig. 14| (Related to Fig. 3).**
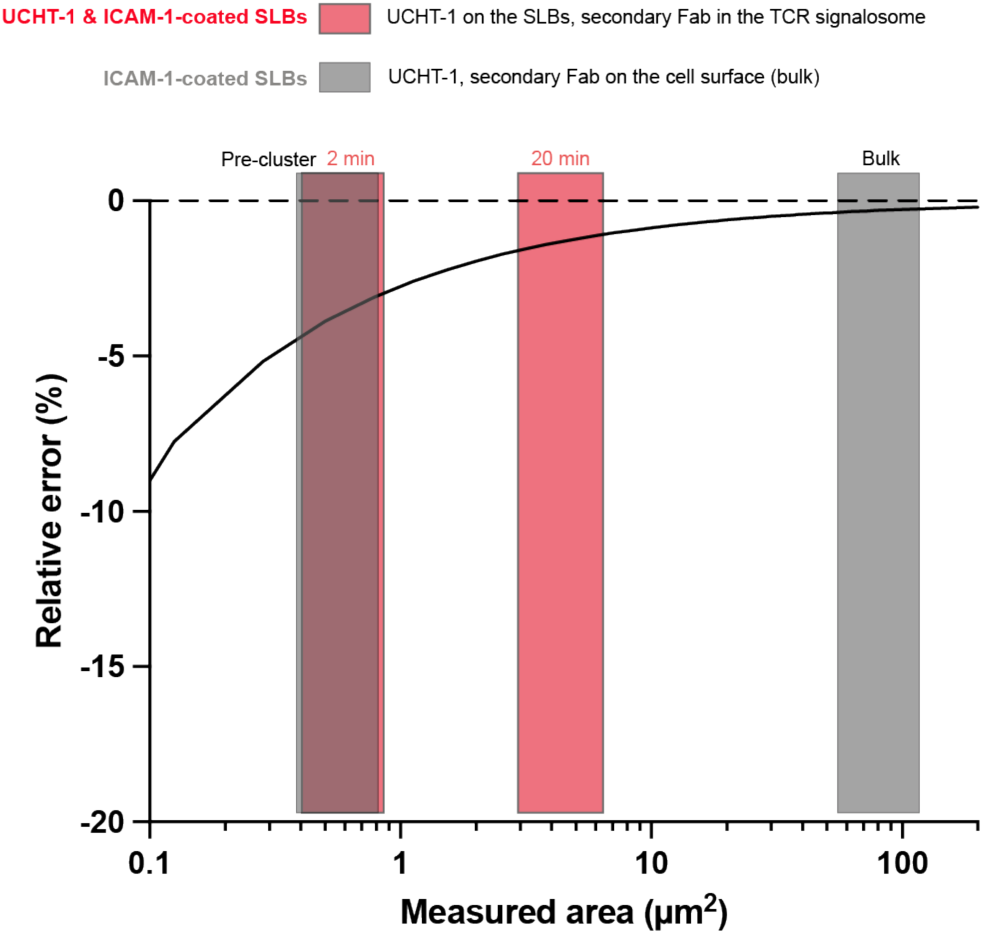
Relative error of molecular quantification of the numbers and densities on the surface or in T cells at different activation states using QuEST.

**Extended Data Fig. 15| (Related to Fig. 4).**
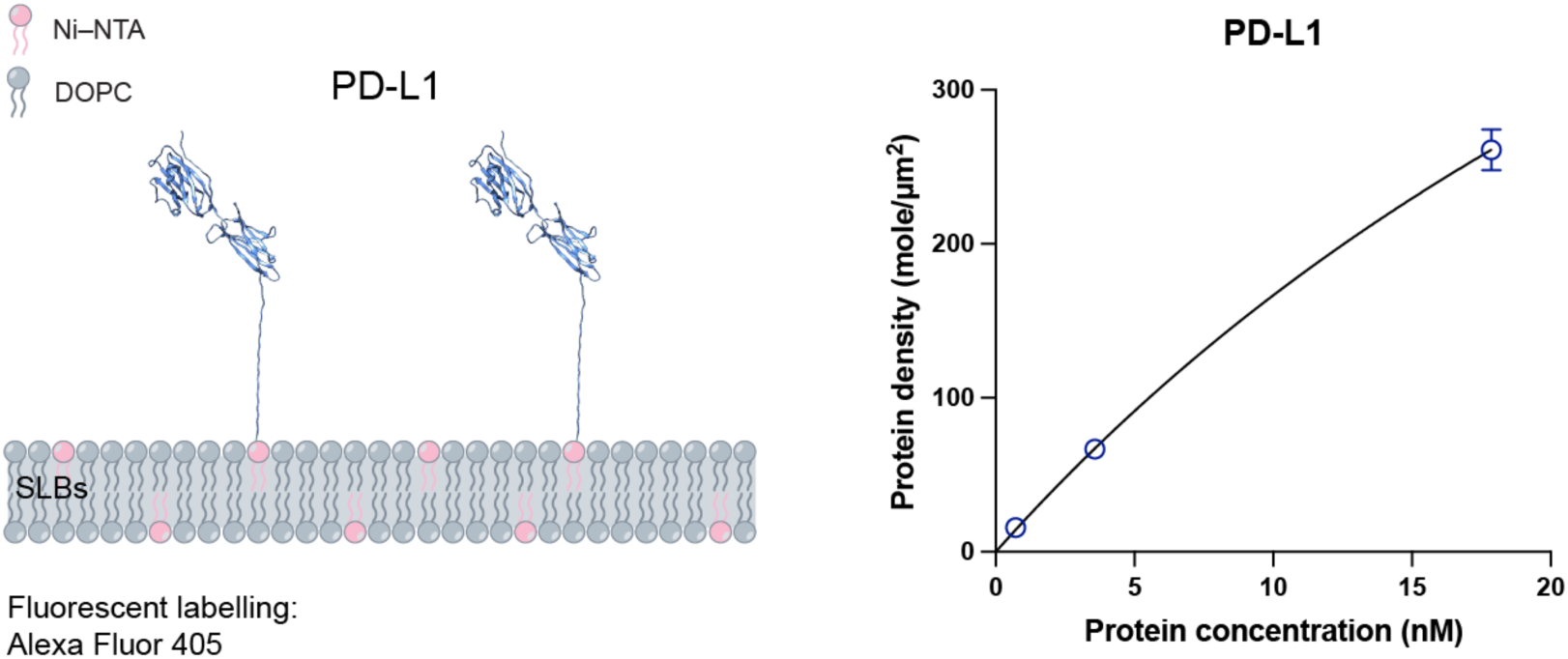
Quantification of PD-L1 density on SLBs using QuEST. Schematic (left) and corresponding quantification results (right) of PD-L1 density on SLBs measured using the QuEST method. The structural model of PD-L1 was generated using AlphaFold3 from the corresponding protein sequence.

**Extended Data Fig. 16| (Related to Fig. 5).**
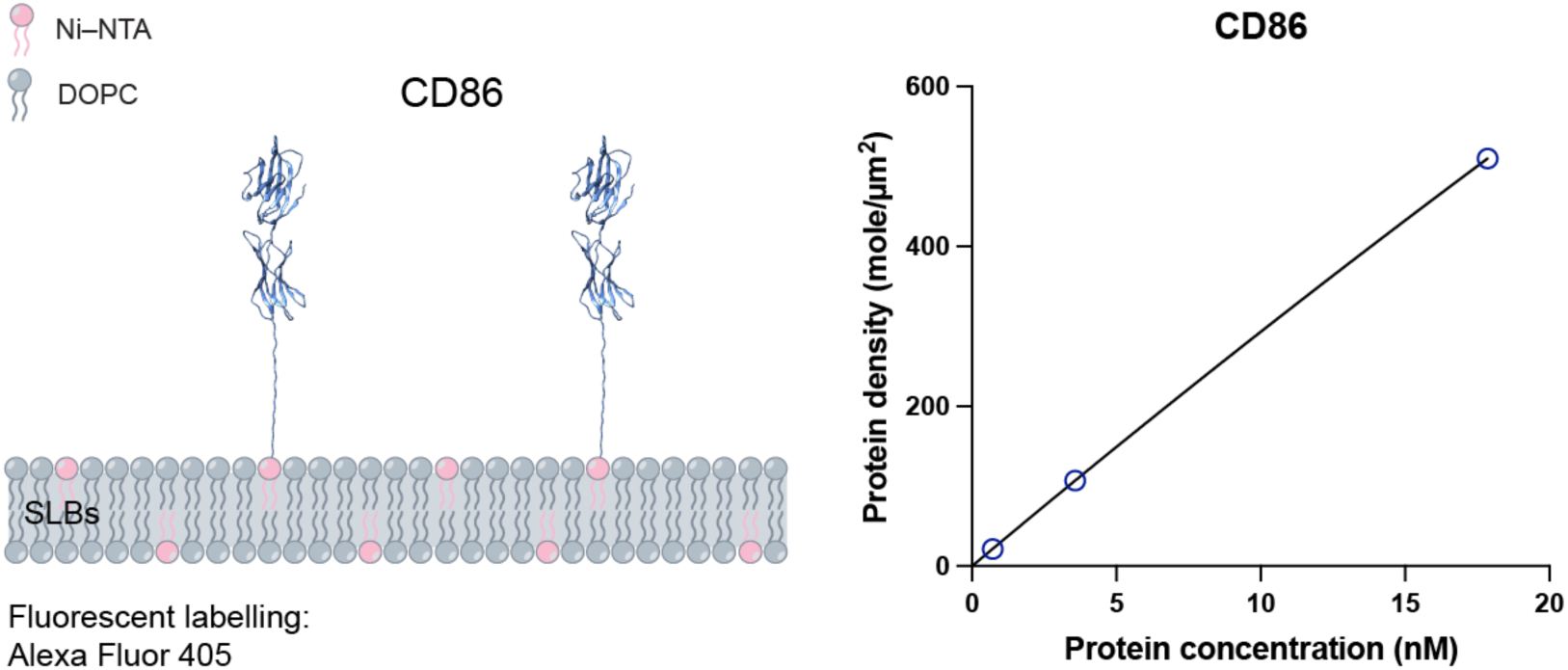
Quantification of CD86 density on SLBs using QuEST. Schematic (left) and corresponding quantification results (right) of CD86 density on SLBs measured using the QuEST method. The structural model of CD86 was generated using AlphaFold3 from the corresponding protein sequence.

## Supplementary Figures and legends

**Supplementary Fig. 1| (Related to Fig. 2).**
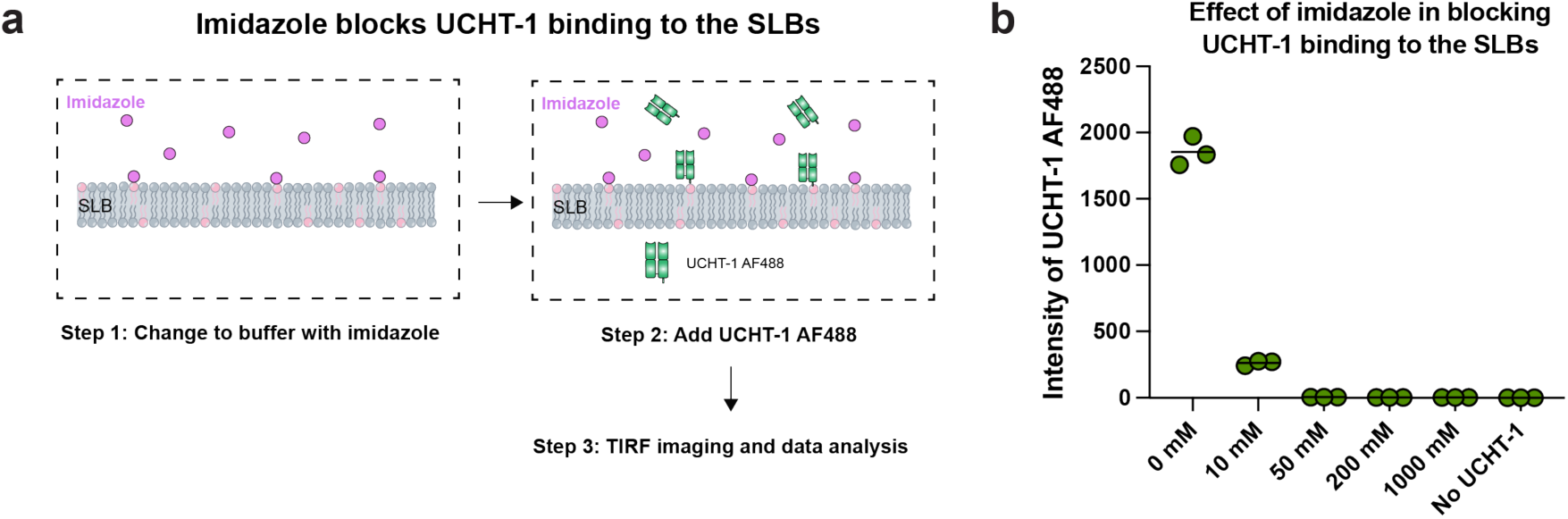
Imidazole blocks UCHT-1 binding to supported lipid bilayers (SLBs). **a**,**b**, Workflow (**a**) and corresponding quantification results (**b**) of the blocking experiment. Imidazole (100 mM) was used in the quantification experiments.

**Supplementary Fig. 2| (Related to Fig. 3).**
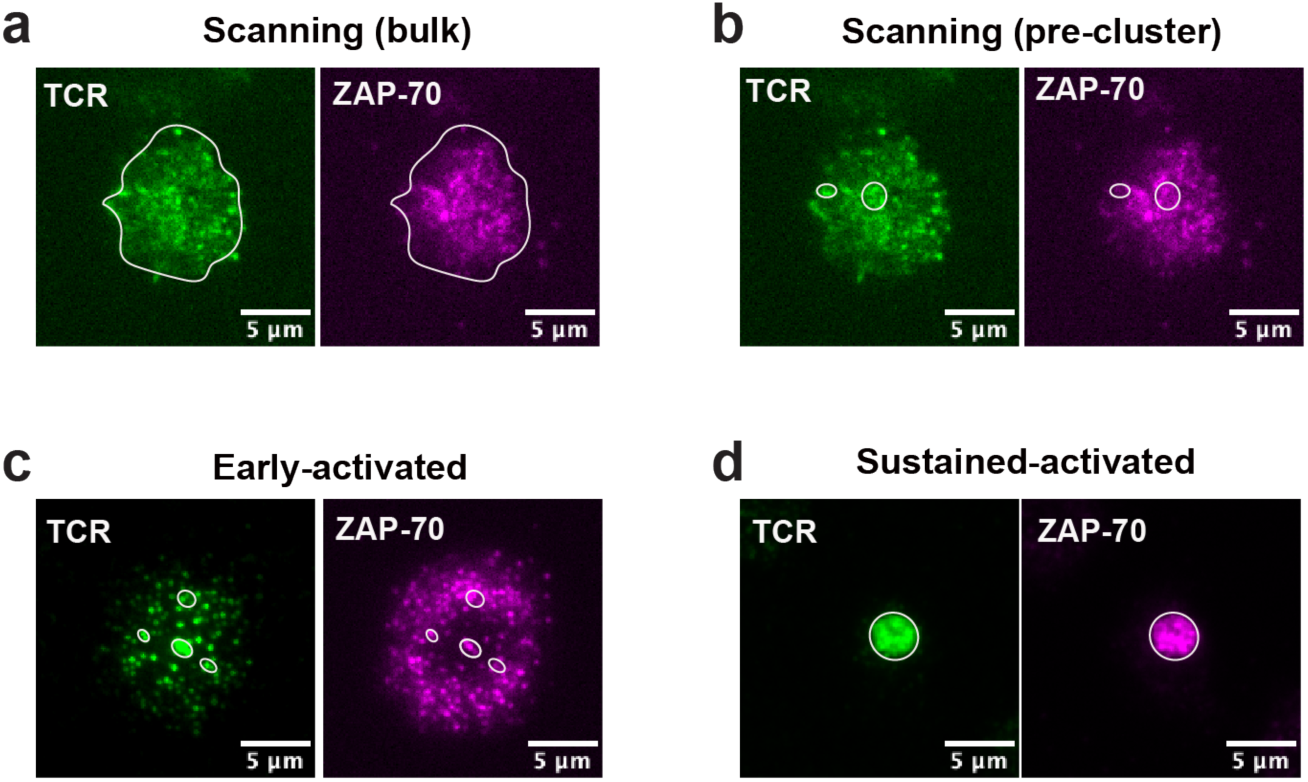
The regions selected for analysis at different stages of T cell activation are indicated by white outlines. **a**, Scanning (bulk) state. **b**, Scanning (pre-cluster) state. **c**, Early-activated state. **d**, Sustained-activated state.

**Supplementary Fig. 3| (Related to Fig. 3).**
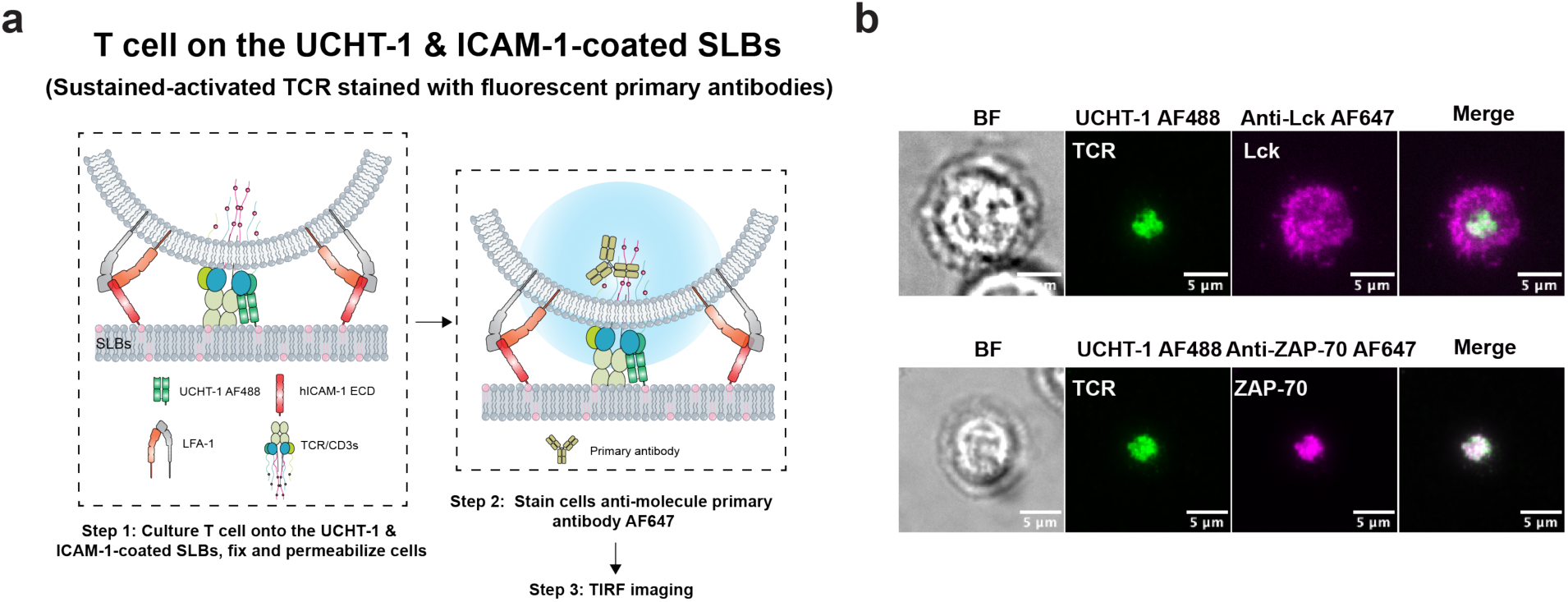
Fluorescent primary antibody staining of Lck or ZAP-70. **a**,**b**, Workflow (**a**) and representative TIRF images (**b**) of Lck or ZAP-70 staining using fluorescently labelled primary antibodies during sustained TCR activation (20 min). Data are representative of two replicates using T cells from two blood donors.

**Supplementary Fig. 4| (Related to Fig. 3).**
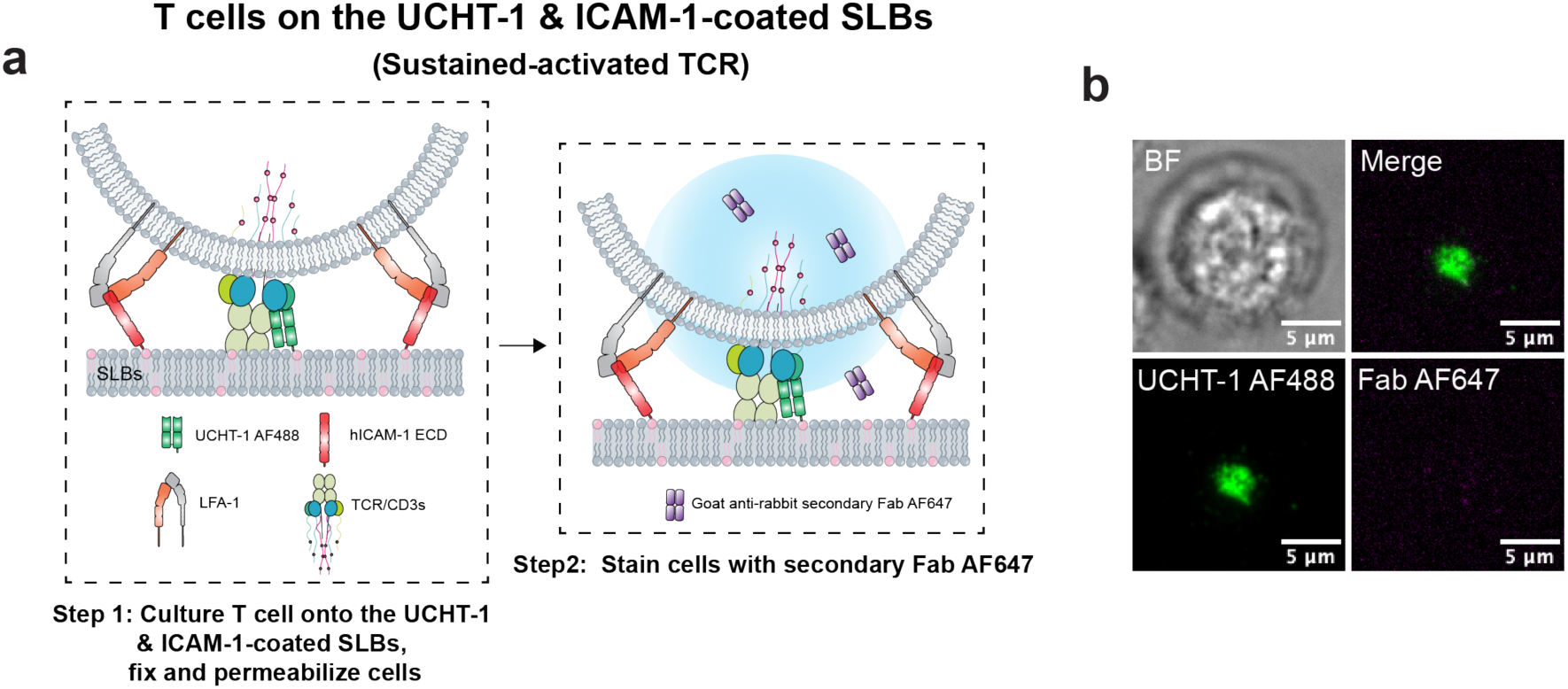
Verification of cross-staining by goat anti-rabbit secondary Fab. **a**,**b**, Workflow (**a**) and representative TIRF images (**b**) assessing cross-staining of goat anti-rabbit secondary Fab to UCHT-1 in T cells upon TCR activation. Data are representative of two replicates using T cells from two blood donors.

## References

1. Grakoui, A. et al. The immunological synapse: a molecular machine controlling T cell activation. Science 285, 221–227 (1999).

2. Monks, C.R., Freiberg, B.A., Kupfer, H., Sciaky, N. & Kupfer, A. Three-dimensional segregation of supramolecular activation clusters in T cells. Nature 395, 82–86 (1998).

3. Werlen, G. & Palmer, E. The T-cell receptor signalosome: a dynamic structure with expanding complexity. Curr. Opin. Immunol. 14, 299–305 (2002).

4. van der Merwe, P.A. & Dushek, O. Mechanisms for T cell receptor triggering. Nat. Rev. Immunol. 11, 47–55 (2011).

5. Campi, G., Varma, R. & Dustin, M.L. Actin and agonist MHC-peptide complex-dependent T cell receptor microclusters as scaffolds for signaling. J. Exp. Med. 202, 1031–1036 (2005).

6. O’Donoghue, G.P., Pielak, R.M., Smoligovets, A.A., Lin, J.J. & Groves, J.T. Direct single molecule measurement of TCR triggering by agonist pMHC in living primary T cells. Elife 2, e00778 (2013).

7. Varma, R., Campi, G., Yokosuka, T., Saito, T. & Dustin, M.L. T cell receptor-proximal signals are sustained in peripheral microclusters and terminated in the central supramolecular activation cluster. Immunity 25, 117–127 (2006).

8. Paludan, S.R., Pradeu, T., Masters, S.L. & Mogensen, T.H. Constitutive immune mechanisms: mediators of host defence and immune regulation. Nat. Rev. Immunol. 21, 137–150 (2021).

9. Sun, L., Su, Y., Jiao, A., Wang, X. & Zhang, B. T cells in health and disease. Signal Transduct. Target Ther. 8, 235 (2023).

10. George, A.J., Stark, J. & Chan, C. Understanding specificity and sensitivity of T-cell recognition. Trends Immunol. 26, 653–659 (2005).

11. Guy, C.S. et al. Distinct TCR signaling pathways drive proliferation and cytokine production in T cells. Nat. Immunol. 14, 262–270 (2013).

12. Gaud, G. et al. CD3zeta ITAMs enable ligand discrimination and antagonism by inhibiting TCR signaling in response to low-affinity peptides. Nat. Immunol. 24, 2121–2134 (2023).

13. Chakraborty, A.K. & Weiss, A. Insights into the initiation of TCR signaling. Nat. Immunol. 15, 798–807 (2014).

14. Voisinne, G. et al. Kinetic proofreading through the multi-step activation of the ZAP70 kinase underlies early T cell ligand discrimination. Nat. Immunol. 23, 1355–1364 (2022).

15. Shao, W. et al. Novel insights into TCR-T cell therapy in solid neoplasms: optimizing adoptive immunotherapy. Exp. Hematol. Oncol. 13, 37 (2024).

16. Lelek, M., et al. Single-molecule localization microscopy. Nat. Rev. Methods Primers 1 (2021).

17. Rust, M.J., Bates, M. & Zhuang, X. Sub-diffraction-limit imaging by stochastic optical reconstruction microscopy (STORM). Nat. Methods 3, 793–795 (2006).

18. Neuman, K.C. & Nagy, A. Single-molecule force spectroscopy: optical tweezers, magnetic tweezers and atomic force microscopy. Nat. Methods 5, 491–505 (2008).

19. Lei, Y. et al. A loosened gating mechanism of RIG-I leads to autoimmune disorders. Nucleic Acids Res. 50, 5850–5863 (2022).

20. Sugiyama, Y., Kawabata, I., Sobue, K. & Okabe, S. Determination of absolute protein numbers in single synapses by a GFP-based calibration technique. Nat. Methods 2, 677–684 (2005).

21. Chiu, C.S., Kartalov, E., Unger, M., Quake, S. & Lester, H.A. Single-molecule measurements calibrate green fluorescent protein surface densities on transparent beads for use with ‘knock-in’ animals and other expression systems. J. Neurosci. Methods 105, 55–63 (2001).

22. Nusser, Z. A new approach to estimate the number, density and variability of receptors at central synapses. Eur. J. Neurosci. 11, 745–752 (1999).

23. Tanaka, J. et al. Number and density of AMPA receptors in single synapses in immature cerebellum. J. Neurosci. 25, 799–807 (2005).

24. Mutch, S.A. et al. Determining the number of specific proteins in cellular compartments by quantitative microscopy. Nat. Protoc. 6, 1953–1968 (2011).

25. Ulbrich, M.H. & Isacoff, E.Y. Subunit counting in membrane-bound proteins. Nat. Methods 4, 319–321 (2007).

26. Jungmann, R. et al. Quantitative super-resolution imaging with qPAINT. Nat. Methods 13, 439–442 (2016).

27. Junghans, V. et al. Hydrodynamic trapping measures the interaction between membrane-associated molecules. Sci. Rep. 8, 12479 (2018).

28. Ben-Sasson, A.J. et al. Design of biologically active binary protein 2D materials. Nature 589, 468–473 (2021).

29. Regan, D., Williams, J., Borri, P. & Langbein, W. Lipid Bilayer Thickness Measured by Quantitative DIC Reveals Phase Transitions and Effects of Substrate Hydrophilicity. Langmuir 35, 13805–13814 (2019).

30. MacDonald, R.I. Characteristics of self-quenching of the fluorescence of lipid-conjugated rhodamine in membranes. J. Biol. Chem. 265, 13533–13539 (1990).

31. Bunnell, S.C. et al. T cell receptor ligation induces the formation of dynamically regulated signaling assemblies. J. Cell. Biol. 158, 1263–1275 (2002).

32. Cassioli, C. et al. Increasing LFA-1 Expression Enhances Immune Synapse Architecture and T Cell Receptor Signaling in Jurkat E6.1 Cells. Front. Cell Dev. Biol. 9, 673446 (2021).

33. Heesink, G. et al. Quantification of Dark Protein Populations in Fluorescent Proteins by Two-Color Coincidence Detection and Nanophotonic Manipulation. J. Phys. Chem. B. 126, 7906–7915 (2022).

34. Axelrod, D. Chapter 7: Total internal reflection fluorescence microscopy. Methods Cell Biol. 89, 169–221 (2008).

35. Crites, T.J. et al. TCR Microclusters pre-exist and contain molecules necessary for TCR signal transduction. J. Immunol. 193, 56–67 (2014).

36. Zhao, Y. et al. cis-B7:CD28 interactions at invaginated synaptic membranes provide CD28 co-stimulation and promote CD8(+) T cell function and anti-tumor immunity. Immunity 56, 1187–1203 e1112 (2023).

37. Goyette, J. et al. Dephosphorylation accelerates the dissociation of ZAP70 from the T cell receptor. Proc. Natl. Acad. Sci. U S A 119 (2022).

38. Morch, A.M. et al. The kinase occupancy of T cell coreceptors reconsidered. Proc. Nat. Acad. Sci. U S A 119, e2213538119 (2022).

39. Chan, A.C., Irving, B.A., Fraser, J.D. & Weiss, A. The zeta chain is associated with a tyrosine kinase and upon T-cell antigen receptor stimulation associates with ZAP-70, a 70-kDa tyrosine phosphoprotein. Proc. Natl. Acad. Sci. U S A 88, 9166–9170 (1991).

40. Zhao, Y. et al. PD-L1:CD80 Cis-Heterodimer Triggers the Co-stimulatory Receptor CD28 While Repressing the Inhibitory PD-1 and CTLA-4 Pathways. Immunity 51, 1059–1073 e1059 (2019).

41. Ishida, Y., Agata, Y., Shibahara, K. & Honjo, T. Induced expression of PD-1, a novel member of the immunoglobulin gene superfamily, upon programmed cell death. EMBO J. 11, 3887–3895 (1992).

42. Okazaki, T. & Honjo, T. PD-1 and PD-1 ligands: from discovery to clinical application. Int. Immunol. 19, 813–824 (2007).

43. Mizuno, R. et al. PD-1 Primarily Targets TCR Signal in the Inhibition of Functional T Cell Activation. Front. Immunol. 10, 630 (2019).

44. Pardoll, D.M. The blockade of immune checkpoints in cancer immunotherapy. Nat. Rev. Cancer. 12, 252–264 (2012).

45. Yokosuka, T. et al. Programmed cell death 1 forms negative costimulatory microclusters that directly inhibit T cell receptor signaling by recruiting phosphatase SHP2. J. Exp. Med. 209, 1201–1217 (2012).

46. Kinnunen, P.C., Humphries, B.A., Luker, G.D., Luker, K.E. & Linderman, J.J. Characterizing heterogeneous single-cell dose responses computationally and experimentally using threshold inhibition surfaces and dose-titration assays. NPJ Syst. Biol. Appl. 10, 42 (2024).

47. Hui, E.F. et al. T cell costimulatory receptor CD28 is a primary target for PD-1-mediated inhibition. Science 355, 1428–1433 (2017).

48. Wherry, E.J. & Kurachi, M. Molecular and cellular insights into T cell exhaustion. Nat. Rev. Immunol. 15, 486–499 (2015).

49. Esensten, J.H., Helou, Y.A., Chopra, G., Weiss, A. & Bluestone, J.A. CD28 Costimulation: From Mechanism to Therapy. Immunity 44, 973–988 (2016).

50. Unternaehrer, J.J., Chow, A., Pypaert, M., Inaba, K. & Mellman, I. The tetraspanin CD9 mediates lateral association of MHC class II molecules on the dendritic cell surface. Proc. Natl. Acad. Sci. U S A 104, 234–239 (2007).

51. Walker, L.S. & Sansom, D.M. The emerging role of CTLA4 as a cell-extrinsic regulator of T cell responses. Nat. Rev. Immunol. 11, 852–863 (2011).

52. Simoncelli, S. et al. Multi-color Molecular Visualization of Signaling Proteins Reveals How C-Terminal Src Kinase Nanoclusters Regulate T Cell Receptor Activation. Cell Rep. 33, 108523 (2020).

53. Coelho, S. et al. Ultraprecise single-molecule localization microscopy enables in situ distance measurements in intact cells. Sci. Adv. 6, eaay8271 (2020).

54. Giannone, G. et al. Dynamic superresolution imaging of endogenous proteins on living cells at ultra-high density. Biophys J. 99, 1303–1310 (2010).

55. Stehr, F., Stein, J., Schueder, F., Schwille, P. & Jungmann, R. Flat-top TIRF illumination boosts DNA-PAINT imaging and quantification. Nat. Commun. 10, 1268 (2019).

56. Kvalvaag, A. et al. Clathrin mediates both internalization and vesicular release of triggered T cell receptor at the immunological synapse. Proc. Natl. Acad. Sci. U S A 120, e2211368120 (2023).

57. Stinchcombe, J.C. et al. Ectocytosis renders T cell receptor signaling self-limiting at the immune synapse. Science 380, 818–823 (2023).

## References

1. Fei, P. et al. Utility of TPP-manufactured biophysical restrictions to probe multiscale cellular dynamics. Bio-Des. and Manuf. 4, 776–789 (2021).

2. Dustin, M.L., Starr, T., Varma, R. & Thomas, V.K. Supported planar bilayers for study of the immunological synapse. Curr. Protoc. Immunol. Chapter 18, 18.13.11–18.13.35. (2007).

3. Heesink, G. et al. Quantification of Dark Protein Populations in Fluorescent Proteins by Two-Color Coincidence Detection and Nanophotonic Manipulation. J. Phys. Chem. B. 126, 7906–7915 (2022).

4. Xu, J., Ma, H. & Liu, Y. Stochastic Optical Reconstruction Microscopy (STORM). Curr. Protoc. Cytom. 81, 12.46.11–12.46.27. (2017).

